# Tempo and mode of morphological evolution are decoupled from latitude in birds

**DOI:** 10.1101/2020.06.25.170795

**Authors:** J. Drury, J. Clavel, J.A. Tobias, J. Rolland, C. Sheard, H. Morlon

**Author notes:** correspondence, Durham University, Department of Biosciences, Stockton Road, Durham DH1 3LE. **Author contributions:** JPD & HM designed the study. JC & JPD developed phylogenetic models. JR contributed tools for analyses of historical biogeography. JAT & CS contributed phenotypic datasets. JPD conducted all analyses. JPD & HM wrote the first draft of the manuscript. All authors contributed to revising the manuscript. **Data accessibility statement:** All datasets used will be submitted to a public repository (e.g., Dryad) upon initial acceptance. All scripts for fitting models are currently available at https://github.com/jonathanpdrury/two_regime_models and will be submitted to the R package RPANDA upon acceptance.

## Abstract

The latitudinal diversity gradient is one of the most striking patterns in nature yet its implications for morphological evolution are poorly understood. In particular, it has been proposed that an increased intensity of species interactions in tropical biota may either promote or constrain trait evolution, but which of these outcomes predominates remains uncertain. Here, we develop tools for fitting phylogenetic models of phenotypic evolution in which the impact of species interactions—namely, competition—can vary across lineages. Deploying these models on a global avian trait dataset to explore differences in trait divergence between tropical and temperate lineages, we find that the effect of latitude on the mode and tempo of morphological evolution is weak and clade- or trait-dependent. Our results indicate that competition does not disproportionately impact morphological evolution in tropical bird families and question the validity of previously reported patterns of slower trait evolution in the tropics.

## Introduction

In many groups of organisms, species richness increases toward lower latitudes—a pattern known as the latitudinal diversity gradient—inspiring generations of biologists to search for the causes and consequences of this gradient [1]. One hypothesis posits that species interactions are stronger in the tropics, and therefore play a more important role in many processes (e.g., diversification) in tropical lineages [2–6] (but see [7]). Previous tests of this ‘biotic interactions hypothesis’ have generally focused on latitudinal gradients in the strength of ecological interactions between predator and prey, herbivore and plant, or pathogen and host [8–11]. Latitudinal gradients in the strength of competition between members of the same trophic level have been less explored, although they have been highlighted as one of the most important research directions for testing the biotic interaction hypothesis [5]. Competition among closely related species, such as those from the same taxonomic family, are often assumed to be strong since their ecological and phenotypic similarity increases the likelihood of competition for access to resources or space [12–16]. Such interactions influence selection on traits that mediate access to resources, influencing trait evolution either by promoting divergence between lineages via character displacement [17,18] or, alternatively, by imposing constraints on geographical range overlap and ecological opportunity, reducing trait diversification as niches fill [19–21].

Whether competition predominantly drives or constrains divergence, the impacts on trait evolution should leave a detectable phylogenetic signature [22–25]. In addition, this signature should be most prevalent in the tropics, where each lineage interacts with far larger numbers of potential competitors. As such, the biotic interactions hypothesis predicts differences between tropical and temperate taxa in the pace of evolution (the ‘tempo’, in the parlance of comparative studies) and/or the processes that drive trait diversification (the ‘mode’). In comparison with the wealth of studies that have investigated latitudinal gradients in rates of species diversification [26–30], relatively few have tested for latitudinal gradients in the dynamics of phenotypic evolution, and have mainly focused on tempo rather than mode. Their results so far suggest a potentially complex relationship between trait diversification and latitude. On the one hand, some studies have found greater divergence between sympatric sister taxa in body mass [31] and in plumage coloration [32] in the tropics, supporting the hypothesis that increased competition at lower latitudes drives character displacement [5]. On the other hand, some studies have found that species attain secondary sympatry after speciation more slowly in tropical regions [33], or that evolutionary rates are lower in the tropics for climatic niches [34], body-size [34,35] or social signalling traits [34,36–39], implying that competition may limit ecological opportunity and therefore constrain trait divergence in tropical regions.

Disentangling these opposing effects is challenging because previous macroecological studies have generally been restricted to either relatively few traits or limited samples of species. In addition, previous studies have been impeded by the lack of suitable methods for detecting the impact of species interactions on trait evolution [40–42], although recent progress has been made in developing such methods for use in standard comparative analyses [20,22,24,43,44]. By incorporating species interactions directly into phylogenetic models of trait evolution, these developments overcome some of the issues faced by phylogenetic and trait approaches for studying community assembly that rely on overly simplistic comparisons to randomly assembled communities [43,45,46]. However, these developments have not yet been deployed in the context of latitudinal sampling and thus the key prediction of a latitudinal gradient in trait diversification has yet to be tested.

Here, we begin by expanding existing phylogenetic models of phenotypic evolution, including models that incorporate competition between species — namely, diversity-dependent models [19,20] and the matching competition model [22,43] — such that the impact of competition can be estimated separately in lineages belonging to different, pre-defined competitive regimes (e.g., tropical and temperate). The models we develop are designed to account for known intraspecific variability and unknown, nuisance measurement error, both of which can strongly bias model support and parameter estimates [47]. In particular, it has been suggested that intraspecific variability is lower in the tropics [48], which could inflate estimates of evolutionary rates in the temperate biome. Next, we conduct a comprehensive test of the biotic interactions hypothesis using these new phylogenetic tools to model the effect of interspecific competition on the tempo and mode of morphological evolution based on seven morphological characters describing variation in body size, bill size and shape, and locomotory strategies sampled from ~9400 species representing more than 100 avian families worldwide. These morphological characters have been demonstrated to predict diet and foraging behaviour in birds [49], indicating that they are well suited as proxies for analysing the dynamics of ecological divergence.

## Results

### Latitudinal variation in mode of phenotypic evolution

We tested whether modes of phenotypic evolution varied with latitude using two types of models. First, we tested whether support for various ‘single-regime’ models that estimate a single set of parameters on the entire avian phylogeny [26] varied according to a clade-level index of tropicality. Second, we developed and used ‘two-regime’ models with distinct sets of parameters for tropical and temperate species and tested whether these latitudinal models were better supported than single-regime models.

Across single-regime fits, we found no evidence for a latitudinal trend in the overall support for any model of phenotypic evolution (Fig. 1a-f, S4 Table), with one exception: there was an increase in model support for the matching competition model in tropical lineages for the locomotion pPC3 (Fig. 1f, S4 Table). Similarly, there was no evidence that the overall support for models incorporating competition is higher in tropical clades (Fig. 1g, S4 Table). Models with latitude (i.e., two-regime models) were not consistently better supported than models without latitude, for any model or trait (S5 Table). Indeed single-regime models were the best fit models across 86% of individual clade-by-trait fits (S7 Fig.).

**Figure 1.**
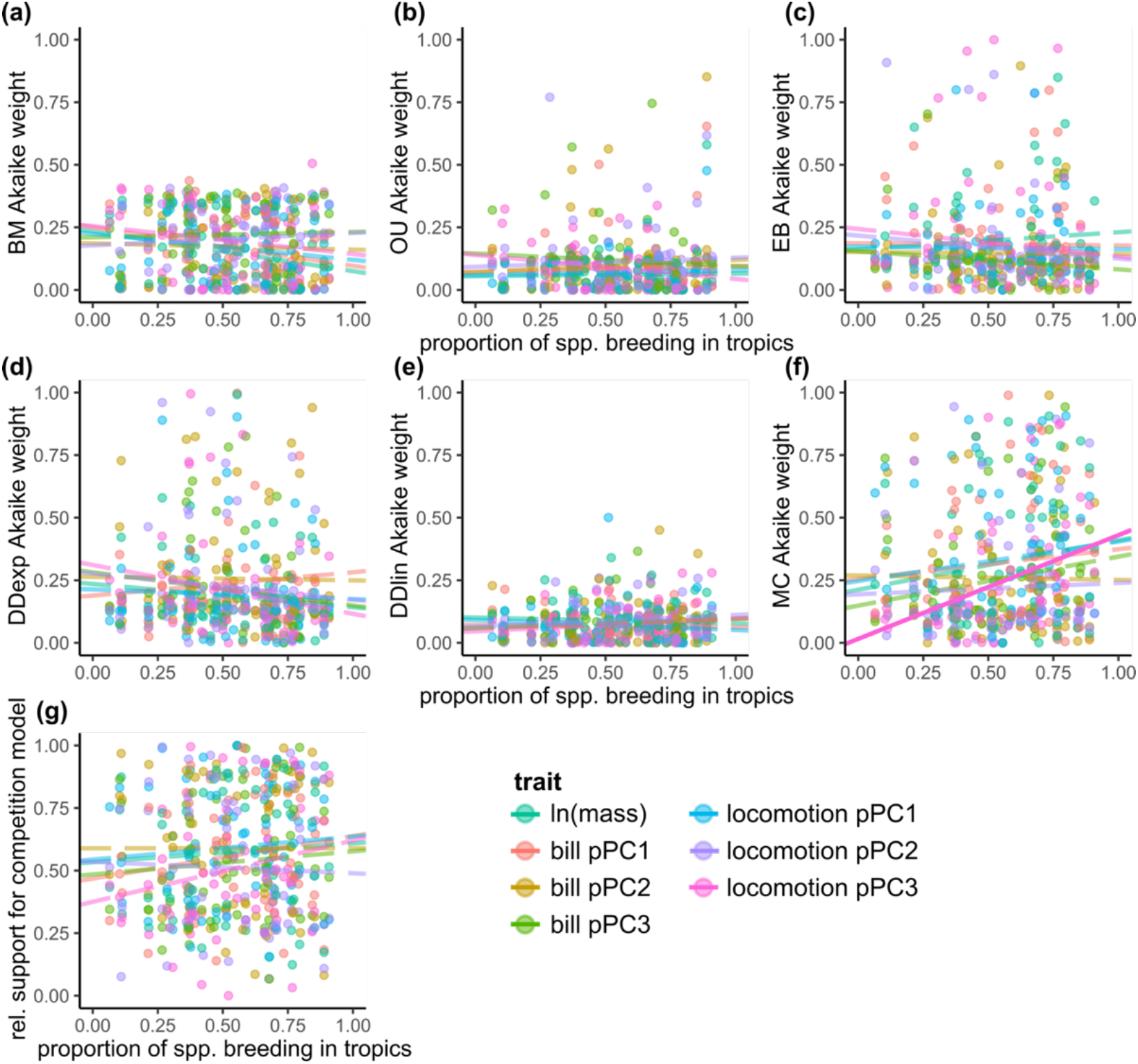
Model support for single-regime models reveal little impact of latitude on the mode of phenotypic evolution in birds (66 clades with ≥ 50 species, with data from 7163 species). There is no relationship between the proportion of taxa in a clade that breed in the tropics and statistical support (measured as the Akaike weight) for **(a)** Brownian motion, **(b)** Ornstein-Uhlenbeck, **(c)** early burst models, **(d)** exponential diversity-dependent models or **(e)** linear diversity-dependent models. In matching competition models **(f)**, there is an increase in model support for locomotion pPC3 (solid line). The relative support for a model incorporating competition does not vary latitudinally for any trait (S4 Table). Each point represents the mean Akaike weight across clade-by-trait fits to stochastic maps of biogeography (i.e., each clade contributes a point for each of seven traits).

### Latitudinal variation in strength of interactions

We found no evidence for a latitudinal trend in the strength of competition estimated from slopes of single-regime diversity-dependent models (Fig. 2c,d, S6 Table). However, the strength of competition estimated from single-regime matching competition models increased in more tropical families for locomotion pPC3 (Fig. 2b, S6 Table). Parameter estimates from two-regime models with competition do not support the biotic interactions hypothesis (Fig. 3b-d)—in most traits, there is no consistent difference between estimates of the impact of competition on tropical and temperate lineages, and in one case (bill pPC2), there is evidence that competition impacts temperate lineages to a larger degree than tropical lineages (Fig. 3b-d, S7 Table). In all cases, there was substantial variation in the fits, and the overall magnitude of differences between tropical and temperate regions was rather small (Fig. 3b-d).

**Figure 2.**
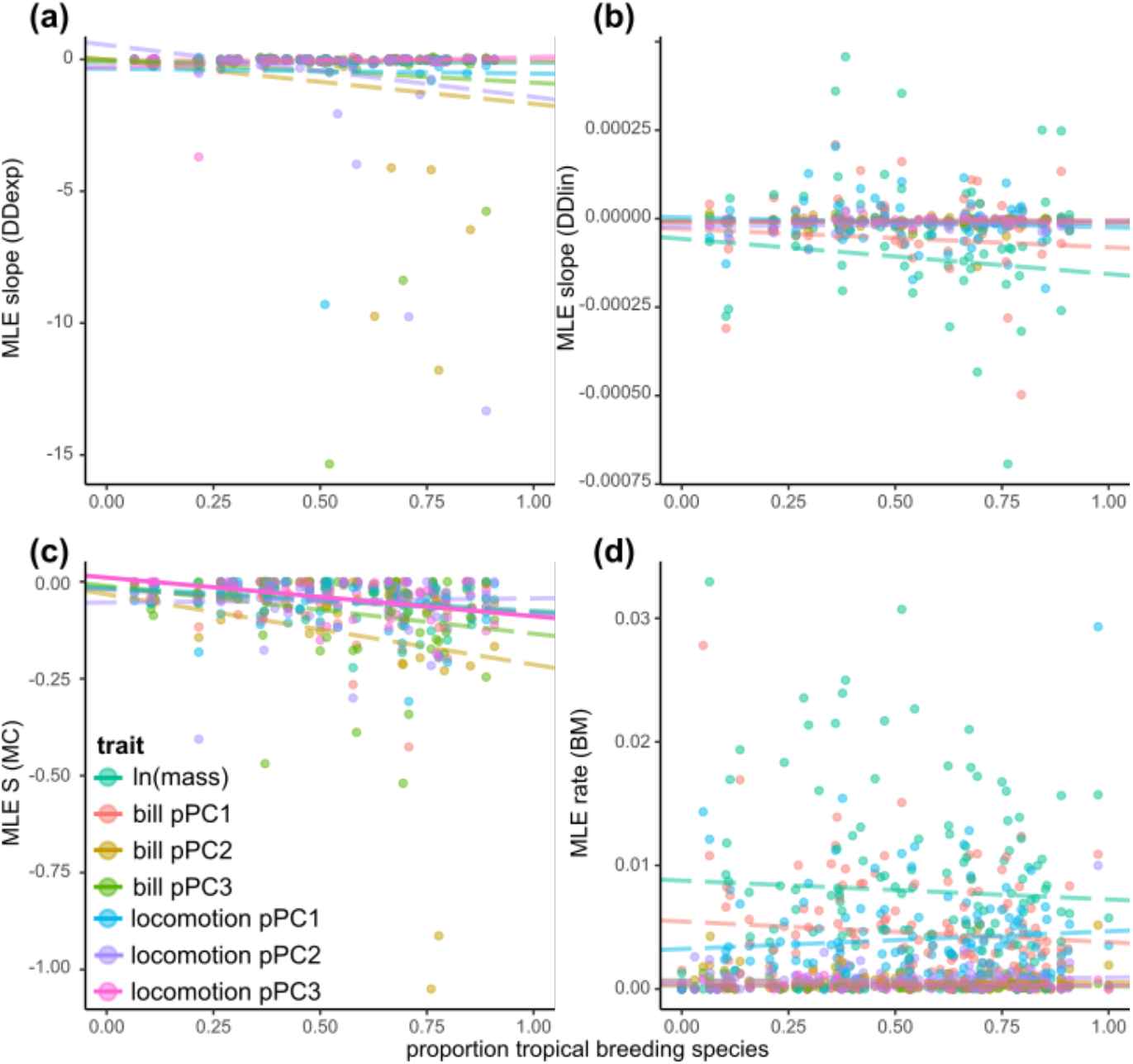
Parameter estimates from single-regime models reveal varying impacts of latitude. There is no latitudinal effect on the effect of competition as measured by the slope of **(a)** exponential diversity-dependent models, or **(b)** linear diversity-dependent models. **(c)** The strength of competition as measured by the S parameter from the matching competition models increases with the index of tropicality (the proportion of species in the clade with exclusively tropical breeding distributions) for locomotion pPC3 but not for other traits. **(d)** There is no relationship between the proportion of taxa in a clade that breed in the tropics and the estimated rate of trait evolution from Brownian motion models. Solid lines represent statistically significant relationships (S6 Table, S13 Table). For (a-c), each point represents the mean across clade-by-trait fits to stochastic maps of biogeography (for all families with at least 50 species), and for (d), each point represents the maximum likelihood estimate for each clade-by-trait fit.

**Figure 3.**
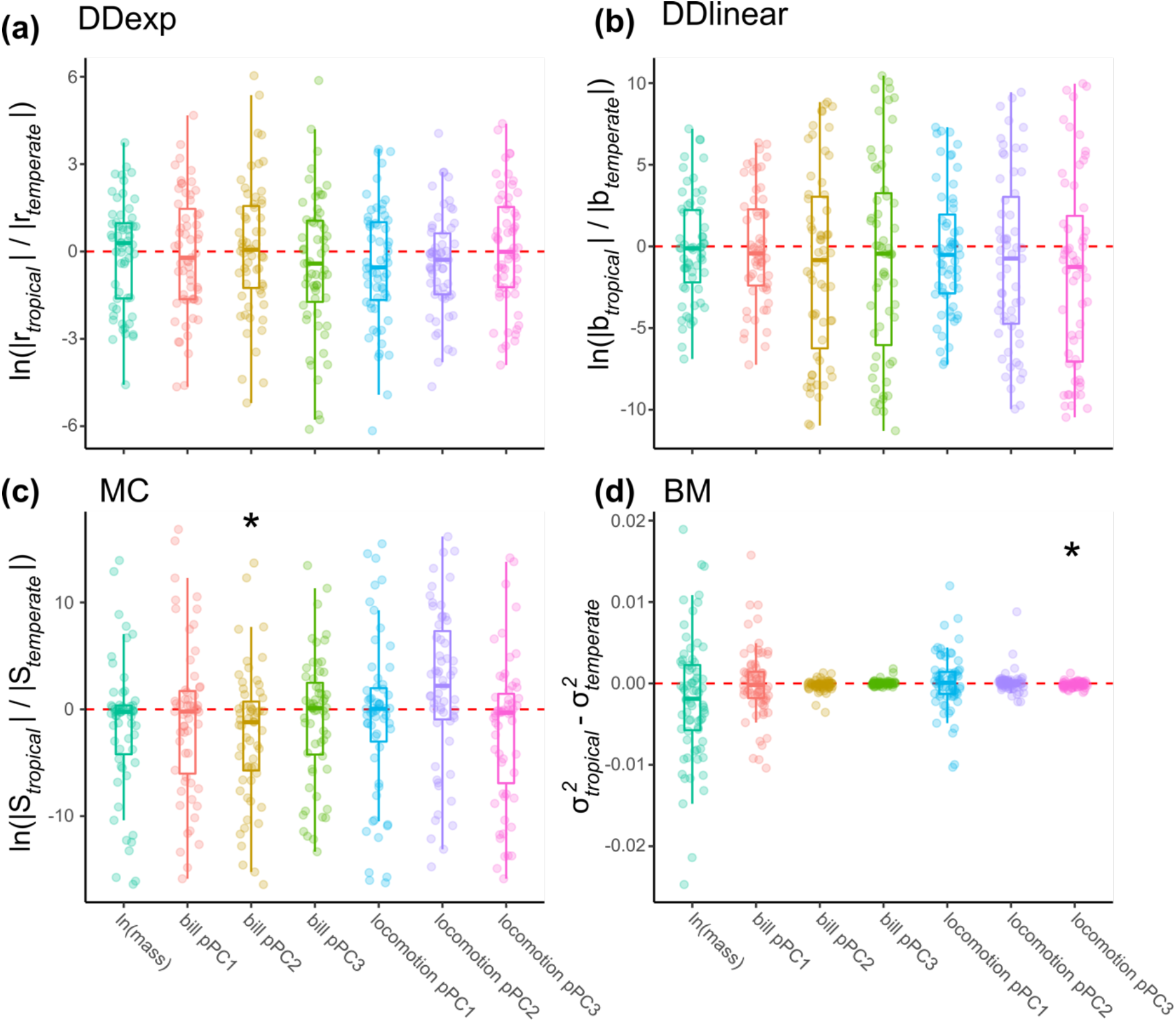
Parameter estimates from two-regime models reveal varying impacts of latitude. Estimates of slopes from **(a)** exponential diversity-dependent models and **(b)** linear diversity dependent models are not consistently different in tropical regions in any trait. **(c)** Matching competition models estimated a decreased strength of competition in the tropics on bill pPC2. **(d)** Estimates of evolutionary rates from Brownian motion models show accelerated rates of locomotion pPC3, but not other functional traits, in temperate regions. Asterisks indicate statistical significance (S7 Table, S14 Table). For **(a-c)**, each point represents the mean across clade-by-trait fits to stochastic maps of biogeography and of tropical/temperate membership (for all families with at least 50 species), and for **(d)**, each point represents the mean across stochastic maps of tropical/temperate membership maximum.

### Impact of assuming continental-scale sympatry

Phylogenetic models of competitively driven trait evolution rely on reconstructions of ancestral ranges to delimit the pool of potential species interaction at each point in the evolutionary history of a clade. Given the scale of our analyses and the computational limits of existing models of ancestral range estimation, we assumed that species occurring on the same continent were able to interact with one another. On average, species in our analyses are sympatric with 50% of clade members at the continental level, although there are differences across continents (mean range 34% - 74%; S3 Fig., S9 Table, S10 Table). Notably, we also found that temperate species are more likely to coexist in sympatry with family members than tropical species (S11 Table). To determine the impact of assuming continental-scale sympatry, we investigated whether we would detect a latitudinal difference in the strength of competition if it existed, even if competition occurrs among only truly sympatric species rather than among all species occurring on the same continent. Simulations examining the impact of the continental-scale sympatry assumption on the statistical power of two-regime MC models demonstrate that, even for relatively small clades, large but biologically plausible latitudinal differences in the strength of competition should be detectable, even when sympatry is overestimated (S8 Fig.). Nevertheless, there is evidence that this assumption can impact the power to detect subtle differences between regions, and for smaller trees, the estimated direction of the difference (S8 Fig.). However, restricting our empirical analyses to large clades (N <u>≥</u> 100), we still find no support for a consistently stronger impact of competition on tropical lineages (S8 Table).

### Latitudinal variation in tempo of phenotypic evolution

Evolutionary rates estimated from single-rate models did not vary according to clade-level index of tropicality (Fig. 2, S9 Fig., S13 Table). Similarly, estimates of rates from latitudinal models were neither consistently lower nor higher in tropical regions (Fig. 3d, S10 Fig, S14 Table). We did find lower rates of locomotion pPC3 (Fig. 3d, S10 Fig., S14 Table) and bill pPC2 evolution in tropical lineages (S10 Fig., S14 Table), but the difference between regions was small and the overall strength of this relationship was weak. Observational error contributed to these patterns: we found a significant negative correlation between observational error and the clade-level index of tropicality for body mass (S11 Fig., S15 Table); we also found that there was a correlation between rates of body mass and locomotion pPC3 evolution in standard single-regime BM models excluding error (S12 Fig., S16 Table), and that the magnitude of the difference between tropical and temperate rates of trait evolution was higher in analyses of two-regime fits excluding error (S12 Fig., S17 Table).

### Predictors of support for models with competition

We found no evidence that territoriality or diet specialization are useful predictors of support for a model with competition (S18 Table). We did, however, find that the maximum proportion of species co-occurring on a continent (i.e., the maximum number of extant lineages on a single continent divided by the total clade size) had a pronounced impact on model selection—clades with a high proportion of lineages occurring on the same continent were more likely to be best-fit by the MC model, whereas clades with a low proportion of co-occurring lineages were more likely to be best-fit by the DD_exp_ model (S13-S14 Figs., S18 Table). In addition, we found that the matching competition model was less likely to be favoured in clades with many members living in single-strata habitats (S18 Table).

## Discussion

Contrary to what would be expected if the strength of competition was stronger in the tropics, we did not find a consistent latitudinal gradient in the dynamics of phenotypic evolution across the entire avian radiation. Using novel methods for examining macroevolutionary signatures of competition, we show that competition between clade members is an important driver of trait evolution but find no evidence that such competition has impacted the dynamics of trait diversification more in the tropics than in temperate regions. This lack of consistent latitudinal effect applied both to the support for specific models of phenotypic evolution and the parameters of these models. Our results contrast with several previous studies that have found a consistent signature of faster rates in the temperate biome [34,36–39,50].

The apparent absence of latitudinal patterns in support of competition models and estimates of competition strength did not arise from overall weak support for competition models, confirming previous findings that competition does leave a detectable signal in comparative, neontological datasets [22–25,51,52]. Indeed, models incorporating species interactions were the best fit models in 25% of clade-by-trait combinations for single-regime fits. In sunbirds (Nectariniidae), for instance, the matching competition model was the best fit model for body mass and two pPC axes describing variation in bill shape, suggesting that competition has driven trait divergence in this diverse clade. In owls (Strigidae), the exponential diversity-dependent model was the best fit model for body mass and several pPC axes describing bill shape and locomotory traits, suggesting that the rate of evolution in owls responds to changing ecological opportunity. The finding that the signature of competition is widespread in birds is echoed by a recent study showing that traits in a similar proportion of clades are best fit by competition models [51].

For both single-regime models and two-regime models, we detected no systematic effect of latitude on the impact of competition on trait diversification. One possible explanation for this is that our approach was highly conservative since we assumed that species occurring on the same continent are likely to interact with one another whereas they may be allopatric (with non-overlapping geographical ranges) or exhibit low levels of syntopy within areas of sympatry [53]. However, previous work [23] and simulations exploring the impacts of assuming competition between potentially allopatric lineages suggest that the MC model is robust to some misspecification of geographic overlap (e.g., allopatric species scored as sympatric). This robustness is likely explained by both the imprint of competition on ancestral, coexisting lineages and a formulation of competition where divergence occurs respective to mean phenotypic values across interacting species (the mean across all species within each continent may be a relatively good proxy for the mean across sympatric species). Nevertheless, the possibility remains that, if differences between regions in the impact of competition are sufficiently small, the two-regime models may not have detected them. In aggregate, however, our results consistently point to a conspicuous absence of a latitudinal gradient in the effect of competition on trait diversification.

One plausible explanation for discrepancies between our results and other studies that examine gradients in the tempo of morphological trait evolution is that our study accounted for observational error. Indeed, we found that overall observational error for body mass increased with latitude; and when we intentionally ignored observational error, Brownian motion models were more likely to pick up faster rates of trait evolution at high latitudes. This result makes sense in the light of previously reported higher trait variance for temperate taxa [48] and a positive correlation between such variance and rate estimates [54]. Our analyses demonstrate that accounting for observational error when testing for latitudinal trends in evolutionary rates is crucial and also suggest that previous analyses overlooking error may have detected spurious latitudinal gradients in trait evolution.

Another potential explanation for the discrepancy between this and previous studies is that many previous studies examined gradients in rapidly evolving plumage and song traits, which may vary latitudinally if sexual or social selection is more pronounced in temperate regions [55]. In contrast, divergence in ecological traits is likely more constrained, as they tend to evolve relatively slowly [56,57].

A third explanation for the discrepancy is that many previous studies used sister-taxa approaches to estimate gradients in trait evolution [34,36,37,50]. Yet, avian sister taxa are younger in temperate regions [33,50], and how these age differences influence rate estimates if trait evolution has proceeded in a non-Brownian fashion is not clear. Analyses on sister taxa of different ages can recover different rates even though these rates are not representative of any process unique to any particular region. For example, given that rates of trait evolution have accelerated toward the present [58]—perhaps because introgression may boost the apparent rates of trait divergence or lead to inaccurate estimates of the timing of splits in young or seemingly young avian lineages [21]—we may expect sister taxa to recover a signature of faster rates in temperate regions (where sister taxa are younger), even if there are no clade-wide latitudinal differences in the overall tempo and mode of evolution.

Within the competition models, the matching competition model was more likely to be chosen as the best-fit model than diversity-dependent models, which is consistent with the notion that competition promotes divergence (e.g., via character displacement [17,18]) more often than it constrains divergence (e.g., via niche saturation [19]) at relatively shallow taxonomic scales [15,42,59]. Although the models we fit are designed to estimate the dynamics of trait evolution, competition can also generate patterns of divergence via its impacts on range dynamics (i.e., ecological sorting) when secondary sympatry is delayed by competitive interactions [21,60,61]. Therefore, although recent evidence suggests that the effects of competitive exclusion on community assembly is distinguishable from the action of character displacement in comparative datasets [25], the possibility remains that the matching competition model may detect a signal of ecological sorting of morphologically distinct lineages [21,62]—a process that is also fundamentally governed by competition—in addition to or instead of evolutionary divergence [25].

In our analyses, we focused within clades, where we would expect competition to be strongest owing to the phenotypic and ecological similarity of recently diverged taxa [16]. Nevertheless, in doing so, we excluded other competitors (e.g., non-family members with similar diets) that impose constraints on niche divergence. Such competitors have been shown to impact rates of trait evolution across clades of birds [54]. Future research could extend our approach by examining the impact of interactions between competitors from a wider diversity of clades.

We found evidence that support for the matching competition model was greater in clades with a higher proportion of lineages occurring on the same continent, suggesting that trait divergence may make coexistence possible [15,18]. The exponential diversity-dependent model, on the other hand, was more likely to be the best-fit model in clades with relatively low levels of continental overlap, which may indicate that in these clades, niche saturation negatively impacts coexistence [63,64]. In addition, we found that model fits on clades with a high proportion of species living in single-strata habitats were less likely to favour the matching competition model, suggesting that opportunity for divergence may be limited in such habitats [65]. These relationships between ecological opportunity, trait evolution, and coexistence highlight the need for models that can jointly estimate the effects of diversification, range dynamics, and trait evolution [25,59]. Such models may identify an impact of competition on processes other than trait evolution, such as competitive exclusion, which may themselves vary latitudinally [21,33].

By including a suite of traits that capture functional variation in niches [49], we were able to identify patterns that would have been highly biased, or that we would have missed, by focusing on one specific trait, in particular body mass. Model support was distributed evenly across different traits, suggesting that the impact of competition varies both across clades and across different functionalities. For instance, while 31% (42/135) of clades exhibit some signature of competition acting on body size evolution in single-regime fits, 68% (92/135) of them exhibit some signature of competition acting on at least one of the seven functional traits (body-size, bill pPC axes and locomotion pPC axes). These results further strengthen the notion that multiple trait axes are necessary to robustly test hypotheses about ecological variation [49,51,66].

We have extended various phylogenetic models of phenotypic evolution, including models with competition, to allow model parameters to vary across lineages (see also [52]) and to account for biogeography and sources of observational error. We then applied them to the case of latitudinal gradients, but they could be used to study other types of geographic (e.g. elevation), ecological (e.g. habitat, diet), behavioural (e.g. migratory strategy) or morphological (e.g. body size) gradients. Studies of gradients in evolutionary rates are often performed using sister-taxa analyses, assuming BM or OU processes [67]. These analyses are useful because they enable quantitative estimates of the impact of continuous gradients on rate parameters. However, by limiting analyses to sister taxa datasets (and therefore ignoring interactions with other coexisting lineages), they are unable to reliably detect signatures of species interactions [68] and so cannot be used to study competition. In addition, these approaches are not well-suited to differentiating between different evolutionary modes. Applying process-based models of phenotypic evolution that incorporate interspecific competition and biogeography allow for such tests of evolutionary hypotheses about the mode of trait evolution across entire clades.

Focusing on competition between closely related species, we did not find support for the biotic interactions hypothesis. Biotic interactions are multifarious; individuals face selective pressures arising from competition, but also from other types of interactions such as predator-prey and host-parasite interactions. Perhaps as a result of this complexity, pinning down clear empirical relationships between latitude and biotic interactions has yielded a complex and often inconsistent set of results [7], with empirical evidence ranging from stronger interactions in the tropics [8,10] to stronger interactions in temperate regions [9]. A challenge for future research on the biotic interactions hypothesis is therefore to more precisely identify the mechanisms that lead to latitudinal gradients in interactions and, consequently, better predict the kinds of interactions that may shape latitudinal gradients in diversification.

## Materials and methods

### Two-regime phylogenetic models of phenotypic evolution

One approach to analyse gradients in phenotypic evolution is to fit phylogenetic models of phenotypic evolution that allow model parameters (e.g., evolutionary rates) to vary across the phylogeny; such models are already available for the simplest models of trait evolution such as Brownian motion (BM) and Ornstein-Uhlenbeck (OU) models [69,70]. To explore effects of species interactions, we developed further extensions to early burst (EB), diversity-dependent (DD) and matching competition (MC) models allowing parameters to be estimated separately in two mutually exclusive groups of lineages in a clade. Generalizing these new models to estimate parameters on more than two groups, or on non-mutually exclusive groups, is straightforward.

We began by developing a two-regime version of the early burst (EB) model in which rates of trait evolution decline according to an exponential function of time passed since the root of the tree [71]. We used this model here to ensure that the diversity-dependent models, which incorporate changes in the number of reconstructed lineages through time, are not erroneously favoured because they accommodate an overall pattern of declining rates through time. To estimate rates of decline separately for mutually exclusive groups, we formulated a two-regime EB model with four parameters (Table 1): z_0_ (the state at the root), σ_0_^2^ (the evolutionary rate parameter at the root of the tree), *r*_A_ (controlling the time dependence on the rate of trait evolution in regime “A”), and *r*_B_ (time dependence in regime “B”). This model can be written as:

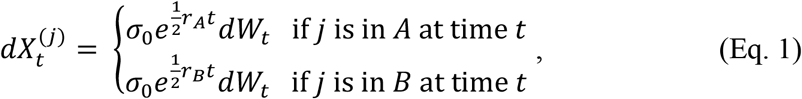

where 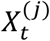 is the trait value of lineage *j* at time *t*, and dW_t_ represents the Brownian motion process (S1 Fig.). This model is the two-regime equivalent of the EB model where *σ*^2^(*t*) = *σ*_0_^2^*e^rt^*; the 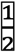 factor in Eq.1 comes from taking the square root of the rate.

**Table 1.**
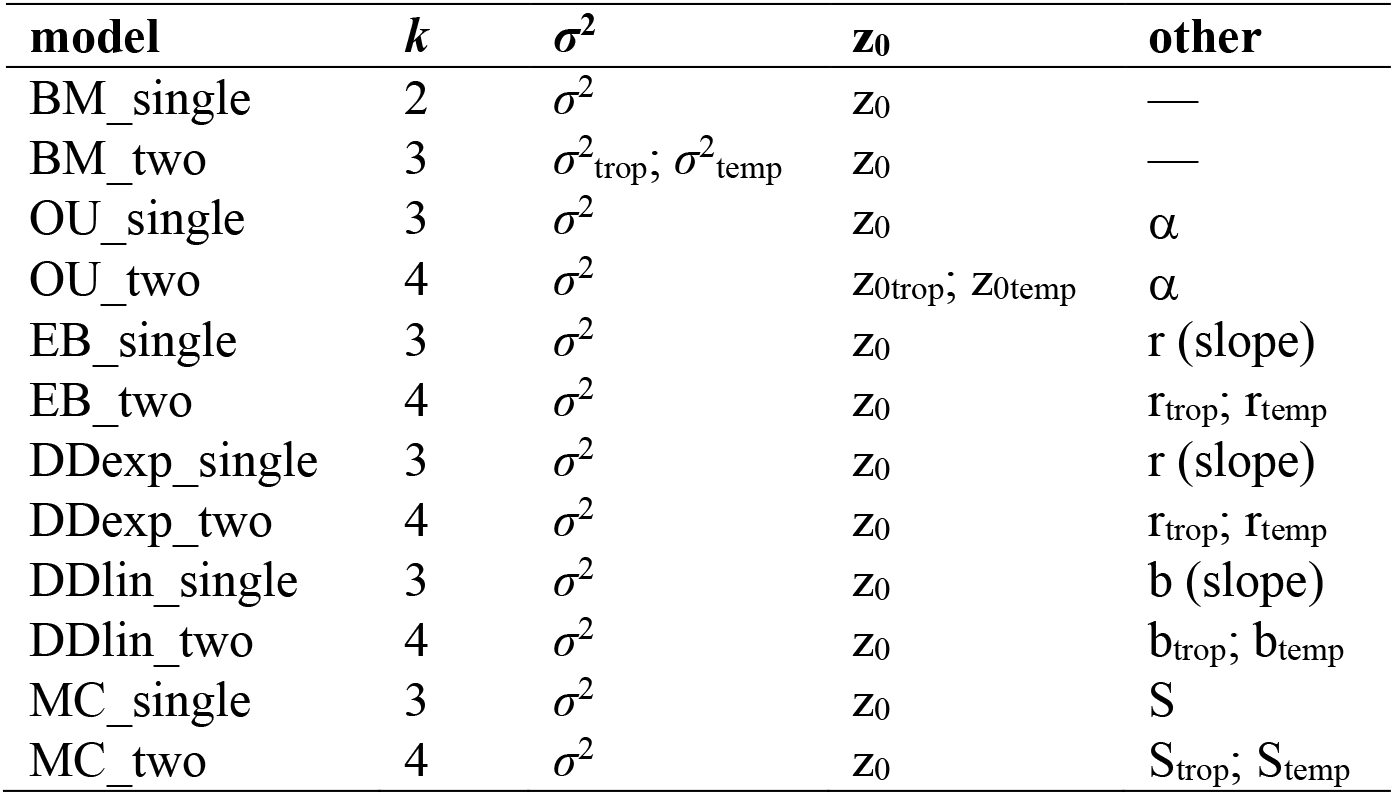
Parameters of models used in analyses. The subscripts ‘trop’ and ‘temp’ in the two-regime versions of each model refer to parameters estimated separately for lineages with exclusively tropical breeding ranges and lineages with breeding ranges that include the temperate region. k indicates the number of free parameters estimated in each model, *σ*^2^ indicates the rate parameter describing the tempo of trait evolution, z_0_ indicates the trait value at the root of the clade, and α describes the strength of the pull toward a stable optimum in the Ornstein-Uhlenbeck model. For descriptions of other parameters, see the main text.

| **model** | **k** | **$\sigma^2$** | **$z_0$** | **other** |
| --- | --- | --- | --- | --- |
| BM_single | 2 | $\sigma^2$ | $z_0$ | — |
| BM_two | 3 | $\sigma^2_{\text{trop}}; \sigma^2_{\text{temp}}$ | $z_0$ | — |
| OU_single | 3 | $\sigma^2$ | $z_0$ | $\alpha$ |
| OU_two | 4 | $\sigma^2$ | $z_{0\text{trop}}; z_{0\text{temp}}$ | $\alpha$ |
| EB_single | 3 | $\sigma^2$ | $z_0$ | r (slope) |
| EB_two | 4 | $\sigma^2$ | $z_0$ | $r_{\text{trop}}; r_{\text{temp}}$ |
| DDexp_single | 3 | $\sigma^2$ | $z_0$ | r (slope) |
| DDexp_two | 4 | $\sigma^2$ | $z_0$ | $r_{\text{trop}}; r_{\text{temp}}$ |
| DDlin_single | 3 | $\sigma^2$ | $z_0$ | b (slope) |
| DDlin_two | 4 | $\sigma^2$ | $z_0$ | $b_{\text{trop}}; b_{\text{temp}}$ |
| MC_single | 3 | $\sigma^2$ | $z_0$ | S |
| MC_two | 4 | $\sigma^2$ | $z_0$ | $S_{\text{trop}}; S_{\text{temp}}$ |

Diversity-dependent (DD) models represent a process where rates of trait evolution respond to changes in ecological opportunity that result from the emergence of related lineages [19,20]. When the slope of these models is negative, this is interpreted as a niche-filling process where rates of trait evolution slow down with the accumulation of lineages. We considered two versions of DD models, with either exponential (DD_exp_) or linear (DD_lin_) dependencies of rates to the number of extant lineages. The two-regime model has four free parameters (Table 1): z_0_ (the state at the root), σ^2^ (the evolutionary rate parameter), *r*_A_ (the dependence of the rate of trait evolution on lineage diversity in regime “A”), and *r*_B_ (diversity dependence in regime “B”). For the exponential case, this model can be written as:

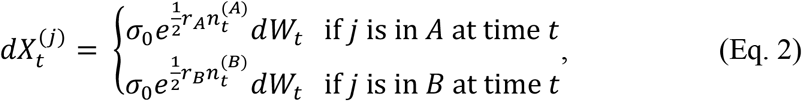

for the exponential case, where 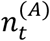 and 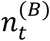 are the number of lineages in regime *A* and *B* at time *t*. This model is the two-regime equivalent of the DD_exp_ model where 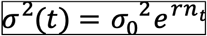; the 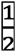 factor in Eq.2 comes from taking the square root of the rate. For the linear case, this can be written as:

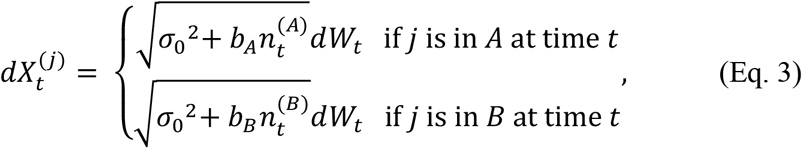

This model is the two-regime equivalent of the DD_lin_ model where *σ*^2^(*t*) = *σ*_0_^2^ + *bn_t_* and *b* denotes the slope in the linear model. Standard DD models ignore whether lineages coexist, yet only those lineages likely to encounter one another in sympatry are able to compete with one another. Thus, we extended our model to incorporate ancestral biogeographic reconstructions to identify which species interactions are possible at any given point in time (i.e., which species co-occur [23]). With biogeography, these become:

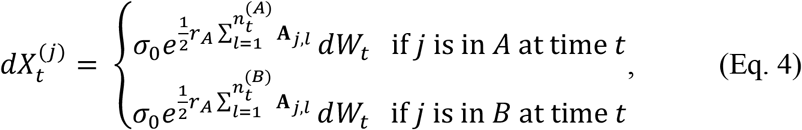

for the exponential case, and:

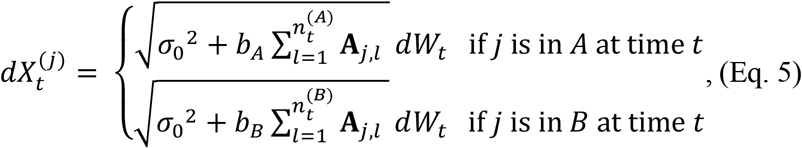

for the linear case, where **A** is a matrix denoting biogeographical overlap, such that **A**_*j,l*_ = 1 if lineages *j* and *l* coexist in sympatry at time *t*, and 0 otherwise (S1 Fig.).

The matching competition (MC) model is a model of competitive divergence [22,43], wherein sympatric lineages are repelled away from one another in trait space. We formulated the two-regime matching competition model, which has four parameters (Table 1): z_0_ (the state at the root), σ^2^ (the evolutionary rate parameter), *S*_A_ (the strength of competition in regime “A”), and *S*_B_ (the strength of competition in regime “B”). This model can be written:

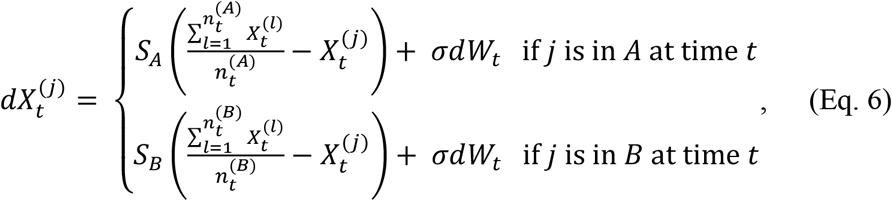

Incorporating biogeography, this becomes:

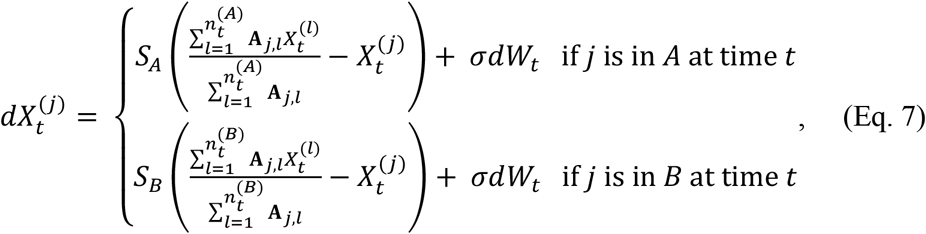

We developed inference tools for fitting the two-regime MC and DD models to comparative trait data, following the numerical integration approach used previously [44,57]. For the EB model, we developed a branch transformation approach similar to the one used in mvMORPH [72]. In all model fits, we incorporated the possibility to account for deviations between measured and modelled mean trait values for each species [73–75] (see S1 Appendix for details). These deviations are of two types: the ‘known’ deviation associated with estimating species means from a finite sample, and the ‘unknown’ deviation linked to intraspecific variability unrelated to the trait model (e.g. instrument errors and phenotypic plasticity). We follow the common practice of lumping these two sources of deviations (often called ‘measurement error’) and referring to them as ‘observational error’. A simulation study demonstrated the reliability of estimates using these tools (S1 Appendix). Functions to simulate and fit these phenotypic models are available in the R package RPANDA (Morlon *et al.* 2016) (note to reviewers: these tools are at https://github.com/jonathanpdrury/two_regime_models and will be submitted to CRAN upon acceptance).

### Phylogeny and trait data

We obtained phylogenies of all available species from birdtree.org [26] and created a maximum clade credibility tree in TreeAnnotator [76] based on 1000 samples from the posterior distribution. Since the MC and DD models require highly sampled clades [43], we used the complete phylogeny including species placed based on taxonomic data [26] and the backbone provided by Hackett et al. [77]. We then extracted trees for each terrestrial (i.e., non-pelagic) family with at least 10 members (*n* = 108). As island species are generally not sympatric with many other members of their families (median latitudinal range of insular taxa = 1.28°, non-insular taxa = 15.27°), we further restricted our analyses to continental taxa, excluding island endemics and species with ranges that are remote from continental land masses. We gathered data on the contemporary ranges of each species from shapefiles [78].

Mass data were compiled from EltonTraits [79] (*n* = 9442). In addition, we used a global dataset based on measurements of live birds and museum specimens [49] to compile six linear morphological measurements: bill length (culmen length), width, and depth (*n* = 9388, mean = 4.5 individuals per species), as well as wing, tarsus, and tail length (*n* = 9393, mean = 5.0 individuals per species). These linear measurements were transformed into phylogenetic principal component (pPC) axes describing functionally relevant variation in bill shape and locomotory strategies (S1 Appendix, S2 Table, S3 Table)

### Biogeographic data and reconstruction

Phylogenetic models that account for species interactions require identifying lineages that are likely to encounter one another [43]. To discretize the contemporary ranges of each species, we classified them as being present or absent in 11 different global regions [80]: Western Palearctic, Eastern Palearctic, Western Nearctic, Eastern Nearctic, Africa, Madagascar, South America, Central America, India, Southeast Asia, and Papua New Guinea/Australia/New Zealand. To assign each species to the global region(s) they occupied, we used several approaches. As a first pass, we used the maximum and minimum longitude and latitude for species’ (non-breeding) ranges. When the rectangle formed by these values fell entirely within the limits of a given global region, we assigned that region as the range for the focal species. Next, for species that did not fall entirely into one region, we compiled observation data from eBird.org [81] to identify all of the regions that a species occupies using country-level observations. Finally, for species whose ranges could not be resolved automatically using these techniques, we manually inspected the ranges.

We incorporated estimates of the presence/absence of each lineage in each range through time using ancestral range estimation under the DEC model of range evolution [82]. We fit DEC models to range data and phylogenies for each family with the R package BioGeoBEARS [82,83]. Since the continents have changed position over the course of the time period of family appearance (clade age range = 12.84 - 71.88 Mya), we ran a stratified analysis with adjacency and dispersal matrices defined for every 10 My time slice [80]. Using the ML parameter estimates for the DEC model, we then created stochastic maps for each family in BioGeoBEARS, each representing a single hypothesis for which ranges each lineage occupied from the root to the tip of the tree.

### Tropical and temperate breeding habitats and reconstruction

To investigate the impact of latitude on trait evolution in two-regime models, we assigned each species to either the ‘tropical’ or ‘temperate’ regime, based on its breeding range (i.e. a species that breeds exclusively in the temperate zones but migrates to the tropics when not breeding is assigned to the temperate zone). We focused on the breeding ranges of all species as they are likely to be the arena of strongest competition over territorial space and food. To do this, we first assigned each species to either ‘tropical’, ‘temperate’, or ‘both’ based on breeding range limits extracted from range data in shapefiles and defining the tropics as the region between −23.437° to 23.437° latitude. We then fit a continuous-time reversible Markov model where transitions between all categories were allowed to occur at different rates, using make.simmap in phytools [84] on the MCC tree. We then used the maximum likelihood transition matrix to create a bank of stochastic maps under this model, each indicating a possible historical reconstruction of tropical vs. temperate habitats through time from the root to tips (S1 Fig.). In each stochastic map, we collapsed the ‘both’ category & the ‘temperate’ category to compare lineages with exclusively tropical ranges to lineages with breeding ranges that include temperate regions. Therefore, our ‘tropical’ category indicates that a species breeds exclusively in the tropics, and our ‘temperate’ category contains all species with breeding ranges that include the temperate zone (S4 Fig.).

We note that this is a relatively simplistic way of categorizing tropical and temperate membership, and we hope that future methods will enable more sophisticated inferences of historical biogeography alongside paleolatitude and/or paleoclimate. However, given the scope of our analyses, and the emerging evidence that many tropical species ranges have shifted over the timescale of this study [85,86], we opted to keep the results of the historical biogeographical inference and the latitudinal-regime reconstruction independent. Future extensions may accommodate the development of more sophisticated paleolatitude models, as well as interactions between various abiotic (e.g., global climate fluctuation [58]) and biotic factors.

### Accounting for uncertainty in historical biogeography and latitude

We accounted for uncertainty in ancestral reconstructions by fitting phenotypic models on at least 20 stochastic maps of ancestral tropical/temperate range membership (for all two-regime models) and/or biogeography (for all models incorporating competition, in both single- and two-regime versions). For the single-regime model fits that included competition (i.e. DD and MC models), we computed model support and parameter estimates as means across fits conducted on stochastic maps of ancestral biogeography. For the two-regime model fits, we computed model support and parameter estimates as means across fits conducted on stochastic maps of ancestral tropical/temperate range membership. For the two-regime model fits with competition, these means also account for variation in estimates of ancestral biogeography (S1 Fig.).

Given the scope of these analyses, we chose to account for uncertainty in the biogeographic reconstructions and in the ancestral reconstruction of tropical/temperate living while keeping the topology fixed under the MCC tree. A previous study with a similar model fitting approach found that results on MCC trees were highly concordant with results fit to trees sampled from the posterior distribution [57]. Moreover, there is no reason, to our knowledge, why basing inferences on the MCC tree would bias conclusions about latitude in any systematic way.

### Latitudinal variation in mode of phenotypic evolution

We tested whether modes of phenotypic evolution varied with latitude in several ways. First, we used ‘single-regime’ models (Table 1), that is, models that estimate a single set of parameters on the entire phylogeny regardless of whether lineages are tropical or temperate. We tested whether support for each of these single-regime models varied according to a clade-level index of tropicality (i.e., the proportion of species in each clade with exclusively tropical breeding ranges). Second, we used our newly developed ‘two-regime’ models (Table 1) with distinct sets of parameters for tropical and temperate species and tested whether these latitudinal models were better supported than models without latitude.

We used maximum likelihood optimization to fit several ‘single-regime’ models of trait evolution to the seven morphological trait values described above. For all families, we fitted a set of six previously described models [43] that include three models (BM, OU, and EB) of independent evolution across lineages, implemented in the R-package mvMORPH [72], and three further models (DD_exp_, DD_lin_, and MC) that incorporate competition and biogeography, implemented in the R-package RPANDA [87]. For details of reconstruction of ancestral biogeography, see Appendix S1. In the diversity-dependent models, the slope parameters can be either positive or negative, meaning that species diversity could itself accelerate trait evolution (positive diversity-dependence), with increasing species richness driving an ever-changing adaptive landscape [4,68]; or, alternatively, increasing species diversity could drive a concomitant decrease in evolutionary rates (negative diversity-dependence), as might be expected if increases in species richness correspond to a decrease in ecological opportunity [88].

In cases where families were too large to fit because of computational limits for the matching competition model (>200 spp., *n* = 13), we identified subclades to which we could fit the full set of models using a slicing algorithm to isolate smaller subtrees within large families. To generate subtrees, we slid from the root of the tree toward the tips, cutting at each small interval (0.1 Myr) until all resulting clades had fewer than 200 tips. We then collected all resulting subclades and fitted the models separately for each subclade with 10 or more species separately, resulting in an additional 28 clades (*n* = 136 total).

In addition to this set of models, we fitted a second version of each of these models where the parameters were estimated separately for lineages with exclusively tropical distributions and lineages with ranges that include the temperate region (i.e., ‘two-regime’ models, S1 Appendix, S2 Fig.), limiting our analyses to clades with trait data for more than 10 lineages in each of temperate and tropical regions (S1 Fig., for details of ancestral reconstruction of tropical and temperate habitats, see S1 Appendix & S4 Fig.). The BM and OU versions of these latitudinal models were fit using the functions mvBM and mvOU in the R package mvMORPH [72], and the latitudinal EB, MC, and DD models were fitted with the newly-developed functions available in RPANDA [87] (note to reviewers: these tools are at https://github.com/jonathanpdrury/two_regime_models and will be submitted to CRAN upon acceptance).

We examined model support in two ways. First, we calculated the Akaike weights of individual models [89], as well as the overall support for any model incorporating species interactions and overall support for any two-regime model. Second, we identified the best-fit model as the model with the lowest small-sample corrected AIC (AICc) value, unless a model with fewer parameters had a ∆AICc value < 2 [89], in which case we considered the simpler model with the next-lowest AICc value to be the best-fitting model.

### Latitudinal variation in strength of interactions and tempo of phenotypic evolution

We tested for latitudinal variation in the effect of species interactions on trait evolution using both our single- and two-regime model fits. With the first class of model, we tested whether parameters that estimate the impact of competition on trait evolution (i.e., the slope parameters of the DD models and the *S* parameter from the MC model) estimated from our single-regime models varied according to the proportion of lineages in each clade that breed exclusively in the tropics. With the second class of models, we tested whether two-regime models estimated a larger impact of competition on trait evolution in tropical than in temperate lineages.

Similarly, we tested whether lineages breeding at low latitudes experience lower or higher rates of morphological evolution compared to temperate lineages using our two types of models. First, we tested whether rates of morphological evolution varied according to the proportion of lineages in each clade that breed exclusively in the tropics. We estimated this rate directly as the *σ*^2^ parameter from the single-regime BM model. For the single-regime EB and DD models, we calculated estimates of evolutionary rates at the present from estimates of the rate at the root and the slope parameters. Second, we compared rates estimated separately for tropical and temperate lineages from the two-regime implementations of the BM, EB, and DD models. We also examined the impact of observational error on rate estimates by fitting single-regime and two-regime BM models without accounting for observational error.

### Examining the potential impact of assuming continental-scale sympatry

Our biogeographical reconstructions add important realism into models of species interactions. Nevertheless, species that occur on the same continent do not necessarily interact with one another. We conducted a simulation analysis to determine how our ability to detect the impact of competition on trait evolution may be impacted by the fact that only a subset of the species occurring in a given continent are actually sympatric.

First, we determined the proportion of species that are sympatric within each continent. We calculated range-wide overlap for all family members that ever coexist on the same continent from BirdLife range maps [78]. We defined sympatry as 20% range overlap according to the Szymkiewicz-Simpson coefficient (i.e., overlap area/min(sp1 area, sp2 area)). We also determined if overall levels of sympatry vary latitudinally; to do so we subset pairs of taxa whose latitudinal means are separated by less than 25° latitude [36] and calculated the midpoint latitude for each pair.

Next, we conducted a simulation study to determine whether competition unfolding between ‘truly’ sympatric species only (i.e., at a level finer than the course continental scale we employed) would systematically impact the fit (i.e., model selection) or performance (i.e., parameter estimation) of the two-regime competition (MC) models for which we used continental-level sympatry (as in the empirical analyses). To do this, we selected three clades spanning the range of tree sizes, each with some traits best-fit by single-regime MC model, but none best-fit by two-regime MC model (Cracidae.0 [N = 50, N_tropical_ = 38, N_temperate_ =12], Nectariniidae.0 [N=122, N_tropical_ = 89, N_temperate_ = 33], Picidae.1 [N=190, N_tropical_ =86, N_temperate_ =104]). For each of these clades, we simulated two biogeographic scenarios reflecting empirical levels of sympatry (see above). In the first, we downsampled the continental biogeography such that 50% of tropical and 50% of temperate taxa that were estimated to occur in the same continent were sympatric (see S1 Appendix for more details). In the second scenario, to reflect the observed latitudinal variation in sympatry, we downsampled the continental biogeography such that 33% of tropical and 50% of temperate taxa that were estimated to occur in the same continent were sympatric (see S1 Appendix for more details).

With these downsampled biogeographic histories, representing hypothetical range overlap that is more realistic than our continental-level assumption of sympatry, we simulated trait evolution under the two-regime matching competition model. For each clade, we used the mean *σ*^2^ value estimated under the single-regime MC model in empirical fits of a trait that was best-fit by the single-regime MC model. We then varied the ratio of the S_tropical_:S_temperate_ within the range of values in other trait-by-clade combinations where the two-regime MC model was the best-fit model (S12 Table). For each clade, parameter combination, and downsampled biogeographic scenario, we simulated 100 datasets, for a total of 3000 simulated datasets. Finally, we fit the same twelve models that were used in empirical analyses. We conducted model selection to identify the best-fit model for each simulated dataset and assessed whether the estimated ln(|S_tropical_|/|S_temperate_|) had the sign expected given the simulated ratio of S_tropical_:S_temperate_.

### Predictors of support for models with competition

To identify factors other than latitude which influence whether models with competition were favoured by model selection, we examined the impact of habitat (the proportion of species in single-strata habitats), territoriality (the proportion of species with strong territoriality), diet specialization (calculated as the Shannon diversity of diets among species in a clade), clade age, clade richness, and the maximum proportion of species co-occurring on a continent.

### Statistical approach

We tested for an impact of the proportion of species in a clade that breed exclusively in the tropics on model support and parameter estimates in single-regime models by conducting phylogenetic generalised least squares using the pgls function in the R package caper [90], estimating phylogenetic signal (*λ*) using maximum likelihood optimization, constraining values to 0 ≤ *λ* ≤ 1. We tested support for the two-regime versions of each model type (BM, OU, EB, DD and MC) across families for a given trait by fitting intercept-only PGLS models with support for latitudinal models as the response variable. We conducted similar analyses to test overall support for latitudinal models across families for each trait and for differences in parameter estimates for tropical and temperate taxa. We found that statistical support for models incorporating competition was relatively rare in small clades (Fig. S6). As this pattern could be related to lower statistical power in smaller datasets [43], we focused all analyses of evolutionary mode (i.e., model support and parameter estimates from models incorporating competition) on clades with at least 50 species (*n* = 66 for single-regime fits, and *n* = 59 for two-regime fits).

For analyses of predictors of support for models with competition, we used the R package MCMCglmm [91] to fit phylogenetic generalised linear mixed models with categorical response variables indicating whether MC or DD_exp_ models were chosen as the best-fit model.

## Acknowledgements

We thank Isaac Overcast, Ignacio Quintero, and other members of the Morlon lab group for helpful comments and discussion and Nick Matzke for assistance generating stochastic maps in BioGeoBEARS. This research was funded by the European Research Council (616419-PANDA to HM) and Natural Environment Research Council (NE/I028068/1 and NE/P004512/1 to JAT).

## Supporting Information Captions

**S1 Appendix.** Supplementary Methods

**S1 Table.** Parameters used for simulations generating datasets used to test two-regime models.

**S2 Table.** Description of morphological variables.

**S3 Table.** Loadings for pPC axes of bill and locomotion measurements.

**S4 Table.** Phylogenetic generalised least-squares (PGLS) models of statistical support as a function of the latitudinal distribution (measured as the proportion of lineages with individuals that breed in tropical regions). Statistical support was measured as the mean Akaike weights of single-regime models (i.e., calculated from pool of single regime models only), and relative support for a model with competition, (defined as the maximum Akaike weight for a model with competition divided by the sum of this value and the maximum Akaike weight for a model without competition [max(MC_wi_, DD_lin_wi_, DD_exp_wi_)/((max(BM_wi_,OU_wi_,EB_wi_)+max(MC_wi_, DD_lin_wi_, DD_exp_wi_))], limiting analyses to clades with ≥ 50 tips (n =66). Values indicated in bold are those that are significant after controlling for multiple testing (α = 0.05/7). λ indicates the maximum likelihood estimate of the phylogenetic signal.

**S5 Table.** Intercept-only PGLS models fit to indices of support for two regimes models for each trait (for cases where N ≥ 50; *n* = 59). The index of relative support for any two-regime model was calculated using max(two regime Akaike weight)/(max(two regime Akaike weight)+max(single regime Akaike weight)); other, model specific indices were calculated using max(two regime Akaike weight for specified model)/ (max(two regime Akaike weight for specified model) + max(single regime Akaike weight for specified model)). For each model, this index was transformed by subtracting 0.5 such that negative estimates indicate support for a single-regime model and positive values equal support for a two-regime model. Values indicated in bold are those that are significant after controlling for multiple testing (α = 0.05/7). For all significant cases, the single-regime version of the model was supported over the two-regime version. λ indicates the maximum likelihood estimate of the phylogenetic signal.

**S6 Table.** PGLS models comparing the observed latitudinal distribution (measured as the proportion of lineages with individuals that breed in tropical regions) of clade-by-trait level fits with the mean maximum likelihood estimates (across fits conducted on a bank of stochastic maps of ancestral biogeography) of the strength of species interactions in single-regime models incorporating competition. All comparisons were conducted on clades with ≥ 50 species (*n* = 66). Note: one outlier was removed from the exponential diversity dependence analysis of bill pPC2. Values indicated in bold are those that are significant after controlling for multiple testing (α = 0.05/7). λ indicates the maximum likelihood estimate of the phylogenetic signal.

**S7 Table.** Intercept-only PGLS models linear regressions fit to tropical/temperate comparisons of maximum likelihood parameter estimates of the strength of species interactions in two-regime models (for cases where N ≥ 50) (*n* = 59) for each trait. For each evolutionary model (a: MC, b: DD_exp_, c: DD_lin_), the mean (across fits conducted on a bank of stochastic maps of ancestral biogeography and stochastic maps of breeding range) of the log-transformed ratio of the absolute value of parameter estimates for tropical taxa to that of temperate taxa (ln(|par_tropical|/|par_temperate|)) was the response variable in the intercept-only PGLS model. Negative estimates, therefore, indicate that the impact of competition is estimated to be higher in temperate regions, whereas positive estimates indicate that competition is higher in the tropics. Values indicated in bold are those that are significant after controlling for multiple testing (α = 0.05/7). λ indicates the maximum likelihood estimate of the phylogenetic signal.

**S8 Table.** Intercept-only PGLS models linear regressions fit to tropical/temperate comparisons of maximum likelihood parameter estimates of the strength of species interactions in two-regime models (for cases where N ≥ 100) (*n* = 34) for each trait. For each evolutionary model (a: MC, b: DD_exp_, c: DD_lin_), the mean (across fits conducted on a bank of stochastic maps of ancestral biogeography and stochastic maps of breeding range) of the log-transformed ratio of the absolute value of parameter estimates for tropical taxa to that of temperate taxa (ln(|par_tropical|/|par_temperate|)) was the response variable in the intercept-only PGLS model. Negative estimates, therefore, indicate that the impact of competition is estimated to be higher in temperate regions, whereas positive estimates indicate that competition is higher in the tropics. Values indicated in bold are those that are significant after controlling for multiple testing (α = 0.05/7). λ indicates the maximum likelihood estimate of the phylogenetic signal.

**S9 Table.** Zero-intercept mixed-effect linear model with a random effect for clade identity fit to the proportion of lineages pairs in each clade that are sympatric in each continent.

**S10 Table.** Intercept-only mixed-effect linear model with a random effect for clade identity fit to the proportion of lineages pairs that are sympatric in each clade.

**S11 Table.** Linear model fit to the proportion of lineages pairs that are sympatric as a function of the absolute value of midpoint latitude for species pairs.

**S12 Table.** Simulation parameters used in simulation study to explore the statistical power of two-regime MC models under realistic levels of sympatry. Values were chosen based on maximum likelihood estimates (MLEs) from single-regime MC models.

**S13 Table.** PGLS analyses of maximum likelihood estimates of evolutionary rates in single-regime model fits (*n* = 135) as a function of the latitudinal distribution (measured as the proportion of lineages with individuals that breed in tropical regions). For diversity-dependent models, parameter estimates are the mean estimates across fits conducted on a bank of stochastic maps of ancestral biogeography. λ indicates the maximum likelihood estimate of the phylogenetic signal.

**S14 Table.** Intercept-only PGLS models fit to the difference between tropical and temperate maximum likelihood parameter estimates of evolutionary rates in two-regime models, fit separately for each trait (*n* = 71 for ln.mass and *n* = 70 for other traits). For DD models, the rate parameter was calculated as the mean comparisons between parameter estimates across fits conducted on a bank of stochastic maps of ancestral biogeography and stochastic maps of breeding range. Note: one outlier was removed from the linear diversity dependence analysis of locomotion pPC2 as it was > 2 orders of magnitude larger than the next largest value. Values indicated in bold are those that are significant after controlling for multiple testing (α = 0.05/7). λ indicates the maximum likelihood estimate of the phylogenetic signal.

**S15 Table.** PGLS models comparing the observed latitudinal distribution (measured as the proportion of lineages with individuals that breed in tropical regions) of clade-by-trait level fits (*n* = 135) with the log-transformed error (calculated as the sum of the maximum likelihood estimated error parameter and the clade-level mean squared standard error) in single-regime Brownian motion models. Values indicated in bold are those that are significant after controlling for multiple testing (α = 0.05/7). λ indicates the maximum likelihood estimate of the phylogenetic signal.

**S16 Table.** PGLS models comparing the observed latitudinal distribution (measured as the proportion of lineages with individuals that breed in tropical regions) of clade-by-trait level fits (*n* = 135) with the maximum likelihood parameter estimates of evolutionary rates in single-regime Brownian motion models that do not account for observational error. Values indicated in bold are those that are significant after controlling for multiple testing (α = 0.05/7). λ indicates the maximum likelihood estimate of the phylogenetic signal.

**S17 Table.** Intercept only PGLS models fit to the mean difference (across stochastic maps of tropical and temperate living) in MLE estimates of tropical and temperate rates (from two-rate BM models that do not account for observational error) (*n* = 71 for log-transformed body mass, 70 for other traits). Values indicated in bold are those that are significant after controlling for multiple testing (α = 0.05/7). λ indicates the maximum likelihood estimate of the phylogenetic signal.

**S18 Table.** The factors predicting which clades support models with competition, as revealed by Phylogenetic Generalised Linear Mixed Models (PGLMMs) fit to single-regime clade-by-trait fits (*n* = 924) with a categorical variable indicating (a) that the matching competition was the modal best fit model (i.e., the most common best fit model across fits conducted on a bank of stochastic maps of ancestral biogeography) (*n* = 166) or (b) that the exponential diversity dependent model was the model best fit model (*n* = 66). The influence of the phylogeny was estimated from the random effect component of the PGLMM—the phylogenetic intraclass correlation coefficient is analogous to the λ parameter (often referred to as ‘phylogenetic signal’) estimated from phylogenetic generalized least squares models [92]. To facilitate parameter exploration, we rescaled all predictor variables using *z*-transformations. We used an uninformative, inverse Wishart distribution as a prior for the random effects, a flat prior for the fixed effects, and fixed the residual variance at 1 [93]. To fit the models, we ran an MCMC chain for at least 5 × 10^5^ generations, recording model results every 100 generations and ignoring the first 5 × 10^3^ generations as burn-in. We fit each model four times and merged the four chains after verifying convergence both visually and using Gelman-Rubin diagnostics in the R-package coda [94,95]. Estimates and credibility intervals are therefore calculated from the pooled posterior distributions. The pMCMC (an MCMC derived *p*-value calculated as two times the proportion of estimates in either the positive or negative portion (whichever is smaller) of the posterior distribution) is presented from one chain.

**S1 Figure.** Illustration of our model-fitting approach for clade-level model fits with different strengths of competition in tropical and temperate regions. We combine a matrix of the presence or absence of each lineage in tropical/temperate regions (‘regime matrix’) with a matrix of biogeography (denoted ‘**A’**) to identify the competitive regime of each lineage and the identity of other lineages with which the focal lineage is able to interact with. Blue and red colours in the lower panel denote correspondence between the formula and the biogeography matrix (**A**) and the regime matrix, respectively.

**S2 Figure.** Results of the simulation study demonstrate the maximum likelihood optimisation returns reliable parameter estimates in two-regime models. **a-d.** exponential time-dependent model **e-h.** exponential diversity-dependent model, **i-l.** linear diversity-dependent model, and **m-p.** matching competition model. In all plots, the red lines denote the parameters used to generate the simulated data.

**S3 Figure.** Results of model selection depicting best fitting models for data simulated under (a) two-regime Brownian motion, (b) two-regime Ornstein-Uhlenbeck, and (c) two-regime Early Burst models across a range of parameter values.

**S4 Figure.** Clade-level distributions of tropical, temperate, and widespread breeding (a) sorted by clade name, (b) sorted by proportion of exclusively tropical breeding species, and (c-d) presented as separate histograms. The number following the family name indicates the subclade within that family (see Methods).

**S5 Figure.** Continental variation in the proportion of species that cooccur in sympatry (defined as 20% range overlap).

**S6 Figure.** Clade size impacts the probability that a model incorporating competition is the modal best-fit model (i.e., the most common best fit model across fits conducted on a bank of stochastic maps of ancestral biogeography and stochastic maps of breeding range).

**S7 Figure.** Best-fit models for each clade-by-trait combination shows that single-regime models generally outperform two-regime models, though some clades (e.g., Meliphagidae, Phasianidae) do tend to support models with latitude across several traits. Shown is the modal best-fit model (i.e., the most common best fit model across fits conducted on a bank of stochastic maps of ancestral biogeography across fits conducted on a bank of stochastic maps of ancestral biogeography and stochastic maps of breeding range). The number following the family name indicates the subclade within that family (see Methods).

**S8 Figure.** Results from simulation analyses exploring the impact of assuming continental level sympatry for three clades. (a-c) Best-fit models for data generated under downsampled biogeographic scenario #1 (i.e., 50% of both tropical and temperate lineages set to allopatric at a continental scale). (d-f) Best-fit models for data generated under downsampled biogeographic scenario #2 (i.e., 50% of temperate lineages and 66.6% of tropical lineages set to allopatric at a continental scale). (g-i) The proportion of simulations for which maximum likelihood estimates of the ratio of competition from the two-regime MC model (i.e., ln(|S_tropical_|/|S_temperate_|)) correctly identify the direction of the difference in the strength of competition.

**S9 Figure.** Evolutionary rates in other single-regime models (a: EB, b: DDexp, c: DDlin) do not vary as a function of the proportion of lineages that breed in the tropics. For diversity-dependent models, parameter estimates are the mean estimates across fits conducted on a bank of stochastic maps of ancestral biogeography.

**S10 Figure.** Differences between rates estimated separately on tropical and temperate taxa in two-regime models (a: EB, b:DDexp, c: DDlin). Shown are the mean comparisons between parameter estimates across fits conducted on a bank of stochastic maps of ancestral biogeography and stochastic maps of breeding range (i.e., tropical or temperate). Asterisks indicate statistical significance.

**S11 Figure.** The relationship between the total error (calculated as the log-transformed sum of the maximum likelihood estimated nuisance error parameter from single-regime Brownian motion models and the clade-level mean squared standard error) and the proportion of tropical breeding lineages in a clade is negative for body mass, but not for other traits. Solid lines represent statistically significant relationships (Table S9).

**S12 Figure.** Brownian motion models of trait evolution fit at a clade level when not accounting for observational error reveal a more pronounced relationship between rate and latitude for several traits **a.** There is a negative relationship between the proportion of taxa in a clade that breed in the tropics and the estimated rate of trait evolution from single-rate Brownian motion models for body mass and locomotion pPC3, but not other traits. Colour of points indicate trait (as in panel b). **b.** Differences between rates estimated separately on tropical and temperate taxa in two-rate Brownian motion models are biased toward faster rates in temperate regions for body mass and locomotion pPC3, but not other traits. Shown are the mean comparisons between parameter estimates across fits conducted on a bank of stochastic maps of ancestral biogeography and stochastic maps of breeding range (i.e., tropical or temperate).

**S13 Figure.** Best-fit ‘single-regime’ models for each clade-by-trait combination show that, while Brownian motion is most often the best model, several clades show evidence of matching competition (e.g., Cotingidae, Formicariidae, Malaconotidae, and Paridae) or diversity dependence (e.g., Strigidae, Fringillidae, Columbidae subclade 2) acting on several traits. Shown is the modal best-fit model across fits conducted on a bank of stochastic maps of ancestral biogeography. The number following the family name indicates the subclade within that family (see Methods).

**S14 Figure.** Best-fit single-regime models (modal best fit across fits conducted on a bank of stochastic maps of ancestral biogeography), plotted as a function of total clade size and the number of species in each clade that occur on the same continent. A) All models, B) Matching competition and exponential diversity-dependent models. Each point represents a clade-by-trait combination (i.e., each clade contributes a point for each of seven traits). In both panels, points are jittered slightly to aid visualization.

