## Appendix S1 for "Tempo and mode of morphological evolution are decoupled from latitude in birds"

S1 Appendix

S1-S18 Tables

S1-S14 Figures

**Appendix S1: Supplementary Methods**

*Parameter estimation in two-regime models*

To verify that the model fitting tools we developed for two-regime

models accurately estimate parameters, we conducted a simulation study for each of the models. In this study, we simulated one hundred trees with 50 tips under a Yule process using the pbtree function in the R package phytools [1]. We then simulated two regimes along the tree, one containing 30 lineages and the other containing 20 lineages, using the make.simmap function in phytools. We then simulated traits from the root to the tips of the phylogenies across a range of parameter values (S1 Table), setting the trait value at the root to 0. Finally, we fit the model for the generating process to each simulated dataset and compared the simulated and estimated parameter values.

In each case, the maximum likelihood estimates were reasonable approximations of the simulated parameter values (S2 Fig.). Importantly, the difference between the parameter values of the two regimes was particularly well estimated (S2 Fig. d,h,l,p). Preliminary analyses demonstrated that the ML estimates from the two-regime MC model were sensitive to the starting values of the optimization process. As a result, for each trait fit in our simulation and empirical analyses, we conducted fits using 5 different starting values and used the results from the fit with the highest likelihood.

*Inferences under two-regime BM, OU, and EB models*

To verify that data generated under two-regime models excluding competition (i.e., BM, OU, and EB models) are not best-fit by models that incorporate competition, we conducted a simulation study for each of the models. We used the same EB simulations as above, and using the same trees and regimes generated above, we simulated BM and OU traits along the phylogenies across a range of parameter values (S1 Table) in mvMORPH [2]. Finally, we fit the model for the generating process to each simulated dataset and conducted model selection as described in the main text.

Perhaps as a result of the relatively small number of lineages in each region, there are parts of parameter space where two-regime models are not better supported than single-regime BM models (S3 Fig). However, with the exception of the performance of single-regime DD_linear_ models on two-regime EB simulations, the two-regime models simulated here rarely lead to erroneous inferring the action of species interactions. Given that DD_linear_ models were very rarely chosen as a best-fit model in our empirical analyses (S7 Fig.), we conclude that our inferences about the role of competition are reliable.

*Phylogenetic principal component analyses*

We constructed two sets of phylogenetic principal component axes [3]: one on the species mean of individually log-transformed values for bill length, height, and width data (bill pPCs), and another on the species mean of individually log-transformed values for wing, tarsus, and tail length (locomotion pPCs) (S2 Table, S3 Table). Each of these axes were constructed on the maximum clade credibility (MCC) bird phylogeny, trimmed to exclude species without data. We acknowledge that using pPCs in comparative analysis of traits involves a level of circularity, since BM evolution is an assumption of these transformations. Nevertheless, using phylogenetic, rather than non-phylogenetic, principal component analyses and conducting analyses across all resulting axes, rather than the first one or two, should help to ameliorate the statistical issues that are known to arise in analyses of trait evolution using PC scores [4].

*Accounting for within-species variation*

Ignoring within-species sources of variation in comparative studies can lead to bias in both parameter estimates and model selection [5–9]. This is, for instance, the case with observational error or uncertainty arising from the estimation of species mean phenotype from a finite sample of individuals and populations. On empirical datasets, observational error variance can be confounded by additional within-species component of variance resulting from instrumental errors, the use of a biased sample for estimating the mean as well as from short-lived causes [e.g., phenotypic plasticity, rapid genetic response to the environment or to fluctuating selection] that are not necessarily captured nor related to the (long-term) process used to model the evolution of the traits on the phylogeny. Holding constant this component of variance across species and assuming that it is independent between species, we can model interspecific trait values by decomposing it as a sum of a phylogenetic, unknown (and independent) intra-specific, and known observational error components:

$$\tilde{\boldsymbol{X}}=z_{0}+\delta+\gamma+\varepsilon$$

Where $\delta\sim\mathcal{N}\left( 0,\sigma_{phylo}^{2}\boldsymbol{D} \right), \gamma\sim\mathcal{N}\left( 0,\sigma_{intra}^{2}\boldsymbol{I} \right),$and $\varepsilon\sim\mathcal{N}\left( 0,\sigma_{error}^{2}\boldsymbol{I} \right)$, are all normally distributed random deviates with scale variance $\sigma_{phylo}^{2}$, $\sigma_{intra}^{2}$ and $\sigma_{error}^{2}$, ***I*** is an identity matrix, and ***D*** is the phylogenetic variance-covariance matrix implied by the evolutionary process used to model **X** in the main text. This decomposition corresponds to a linear mixed model [5] where the variance components $\sigma_{phylo}^{2}$ and $\sigma_{intra}^{2}$ are unknown and have to be estimated during the model fit while $\sigma_{error}^{2}$ is the known sampling variance estimated from multiple individual measurements per species (i.e., the squared standard errors). Given that the evolutionary processes considered here are all Gaussian and it is assumed that the intraspecific components of variance are normally distributed, estimation of all the parameters is done by maximizing the multivariate normal likelihood as generally done in comparative studies [2,10,11].

In our model fits, we input $\sigma_{error}^{2}$ for each of the traits. To calculate these values (i.e., the squared standard errors) for pPC scores, we projected individual, log-transformed measurements [12] into pPC space [13], of which there were on average 4-5 per species (see *Materials and methods*), and then calculated the species-level standard error directly on these values. For body mass, we obtained mean values from EltonTraits [14], using source data [15] where possible to calculate the standard error. When source data were not available, we calculated a standard deviation from the range of mass values when standard deviations were not reported [16] or using the averaged standard deviation across species when a sample size and mean were the only values available or for cases where measurements were only made on a single individual [6]. We then transformed standard error estimates to the log scale [17].

*Examining the potential impact of assuming continental-scale sympatry: further details*

To generate downsampled biogeographies, we first trimmed the clade level phylogeny into two separate trees (one for tropical lineages and one for temperate lineages). On each tree, we generated a bank of 100 stochastic maps for each clade simulating the evolution of a categorical trait meant to represent separate geographic regions within continents, using make.simmap in phytools [1] and an Mk model using rTraitDisc in ape [18]. This categorical trait had either two states in each of the tropical and temperate region (for scenario 1) or two states in the temperate region and three in the tropics (for scenario 2). We then joined the simulated traits across lineages to create a trait with 4 states (for scenario 1) or 5 states (for scenario 2). Finally, we merged the matrices indicating continental-scale sympatry (i.e., the ones used in our empirical analyses) with these simulated matrices such that any two lineages that are not in the same simulated state were set to ‘allopatric’ (i.e., $\mathbf{A}_{j,l}$ = 0) in the original biogeography matrix. In scenario 1, this meant that 50% of lineages that were treated as sympatric in our empirical analyses because they occurred on the same continent were set to be allopatric. In scenario 2, this meant that 50% of temperate lineages and 66.67% of tropical lineages that were treated as sympatric in our empirical analyses because they occurred on the same continent were set to be allopatric.

We then simulated traits under the parameter values in S8 Table across the banks of stochastic maps of downsampled biogeographies using the function simulateTipData in RPANDA [19].

**S1 Table.** Parameters used for simulations generating datasets used to test two-regime models.

| **model** | ***σ*^2^ (tips)** | **regime 1 parameter** | **regime 2 parameter** |
| --- | --- | --- | --- |
| *TD-exponential (EB)* | 0.01 | -2.5 | -1 |
|  |  | -2.5 | -2.5 |
|  |  | -2.5 | -5 |
|  |  | -1 | -2.5 |
|  |  | -5 | -2.5 |
| *DD-linear* | 0.2 | -0.005 | 0.0025 |
|  |  | -0.005 | 0 |
|  |  | -0.005 | -0.0025 |
|  |  | -0.005 | -0.005 |
|  |  | -0.005 | -0.001 |
|  |  | 0.005 | -0.005 |
|  |  | 0 | -0.005 |
|  |  | -0.0025 | -0.005 |
|  |  | -0.005 | -0.005 |
|  |  | -0.001 | -0.005 |
| *DD-exponential* | 0.01 | -0.05 | 0.05 |
|  |  | -0.05 | 0 |
|  |  | -0.05 | -0.025 |
|  |  | -0.05 | -0.05 |
|  |  | -0.05 | -0.1 |
|  |  | 0.05 | -0.05 |
|  |  | 0 | -0.05 |
|  |  | -0.025 | -0.05 |
|  |  | -0.05 | -0.05 |
|  |  | -0.1 | -0.05 |
| *MC* | 0.05 | -0.5 | 0 |
|  |  | -0.5 | -0.25 |
|  |  | -0.5 | -0.5 |
|  |  | -0.5 | -1 |
|  |  | 0 | -0.5 |
|  |  | -0.25 | -0.5 |
|  |  | -0.5 | -0.5 |
|  |  | -1 | -0.5 |
| *BM* |  | 1 | 0.5 |
|  |  | 2 | 0.5 |
|  |  | 0.5 | 0.5 |
|  |  | 0.5 | 1 |
|  |  | 0.5 | 2 |
| *OU* | 0.5 (*α* = 2) | 0 | 1 |
|  | 0.5 (*α* = 2) | 0 | 2 |
|  | 0.5 (*α* = 0.5) | 0 | 1 |
|  | 0.5 (*α* = 0.5) | 0 | 2 |

**S2 Table.** Description of morphological variables.

| **trait** | **description** |
| --- | --- |
| Body mass | ln-transformed body mass |
| Bill pPC1 | High scores = small overall beak size (highest score is *Tachornis phoenicobia*; lowest is *Buceros rhinoceros*) |
| Bill pPC2 | High scores = shape of beak is relatively long and thin (highest score is *Ensifera ensifera*; lowest is *Glaucidium mooreorum*) |
| Bill pPC3 | High scores = shape of beak is relatively narrow (highest score is *Chlorostilbon olivaresi*; lowest is *Caprimulgus maculosus*) |
| Locomotion pPC1 | High scores = small overall size (highest score is *Chaetocercus berlepschi*; lowest is *Leptoptilos dubius*) |
| Locomotion pPC2 | High scores = relatively long tail (highest score is *Eupetomena macroura*; lowest is *Grus americana*) |
| Locomotion pPC3 | High scores = relatively long wings and short tarsi (highest is *Panyptila sanctihieronymi*; lowest is *Stipiturus malachurus*) |

**S3 Table.** Loadings for pPC axes of bill and locomotion measurements.

| **trait group** | **trait** | **pPC1** | **pPC2** | **pPC3** |
| --- | --- | --- | --- | --- |
| bill | ln(culmen length) | -0.86 | 0.51 | -0.07 |
|  | ln(bill width) | -0.89 | -0.33 | -0.3 |
|  | ln(bill depth) | -0.92 | -0.17 | 0.34 |
| locomotion | ln(wing length) | -0.83 | -0.28 | 0.48 |
|  | ln(tarsus length) | -0.78 | -0.56 | -0.28 |
|  | ln(tail length) | -0.92 | 0.38 | -0.07 |

**S4 Table.** Phylogenetic generalised least-squares (PGLS) models of statistical support as a function of the latitudinal distribution (measured as the proportion of lineages with individuals that breed in tropical regions). Statistical support was measured as the mean Akaike weights of single-regime models (i.e., calculated from pool of single regime models only), and relative support for a model with competition, (defined as the maximum Akaike weight for a model with competition divided by the sum of this value and the maximum Akaike weight for a model without competition [max(MC_wi_, DD_lin­_wi_, DD_exp_wi_)/((max(BM_wi_,OU_wi_,EB_wi_)+max(MC_wi_, DD_lin­_wi_, DD_exp_wi_))], limiting analyses to clades with $\geq$ 50 tips (n =66). Values indicated in bold are those that are significant after controlling for multiple testing (α = 0.05/7). λ indicates the maximum likelihood estimate of the phylogenetic signal.

| **term** | **trait** | **estimate** | **std. error** | **t-value** | **p-value** | **λ** |
| --- | --- | --- | --- | --- | --- | --- |
| Akaike weight: BM | ln(mass) | -0.16 | 0.07 | -2.15 | 0.04 | 0 |
|  | bill pPC1 | -0.15 | 0.07 | -2.12 | 0.04 | 0 |
|  | bill pPC2 | -0.02 | 0.08 | -0.33 | 0.74 | 0 |
|  | bill pPC3 | 0.02 | 0.07 | 0.30 | 0.77 | 0 |
|  | locomotion pPC1 | -0.10 | 0.07 | -1.41 | 0.16 | 0 |
|  | locomotion pPC2 | 0.05 | 0.07 | 0.66 | 0.51 | 0 |
|  | locomotion pPC3 | -0.11 | 0.08 | -1.36 | 0.18 | 0 |
| Akaike weight: OU | ln(mass) | 0.01 | 0.05 | 0.21 | 0.83 | 0 |
|  | bill pPC1 | 0.02 | 0.07 | 0.34 | 0.74 | 0 |
|  | bill pPC2 | 0.05 | 0.08 | 0.61 | 0.54 | 0 |
|  | bill pPC3 | -0.05 | 0.07 | -0.71 | 0.48 | 0 |
|  | locomotion pPC1 | 0.02 | 0.04 | 0.44 | 0.66 | 0 |
|  | locomotion pPC2 | 0.04 | 0.08 | 0.48 | 0.63 | 0 |
|  | locomotion pPC3 | -0.1 | 0.05 | -2.09 | 0.04 | 0 |
| Akaike weight: EB | ln(mass) | 0.07 | 0.1 | 0.7 | 0.49 | 0 |
|  | bill pPC1 | -0.01 | 0.09 | -0.1 | 0.92 | 0 |
|  | bill pPC2 | -0.01 | 0.09 | -0.09 | 0.93 | 0.68 |
|  | bill pPC3 | -0.07 | 0.06 | -1.13 | 0.26 | 0 |
|  | locomotion pPC1 | -0.01 | 0.08 | -0.13 | 0.9 | 0 |
|  | locomotion pPC2 | -0.09 | 0.11 | -0.84 | 0.41 | 0.42 |
|  | locomotion pPC3 | -0.11 | 0.13 | -0.86 | 0.39 | 0.8 |
| Akaike weight: DDexp | ln(mass) | -0.09 | 0.1 | -0.97 | 0.33 | 0 |
|  | bill pPC1 | 0.09 | 0.09 | 1.07 | 0.29 | 1 |
|  | bill pPC2 | -0.01 | 0.13 | -0.11 | 0.91 | 0 |
|  | bill pPC3 | -0.13 | 0.1 | -1.32 | 0.19 | 0 |
|  | locomotion pPC1 | -0.04 | 0.11 | -0.36 | 0.72 | 0 |
|  | locomotion pPC2 | -0.1 | 0.12 | -0.9 | 0.37 | 0 |
|  | locomotion pPC3 | -0.19 | 0.11 | -1.78 | 0.08 | 0 |
| Akaike weight: DDlin | ln(mass) | -0.02 | 0.04 | -0.56 | 0.58 | 0 |
|  | bill pPC1 | -0.01 | 0.04 | -0.27 | 0.79 | 0 |
|  | bill pPC2 | 0.04 | 0.05 | 0.77 | 0.45 | 0 |
|  | bill pPC3 | 0.04 | 0.04 | 1.02 | 0.31 | 0 |
|  | locomotion pPC1 | -0.04 | 0.04 | -0.96 | 0.34 | 0.84 |
|  | locomotion pPC2 | 0.05 | 0.04 | 1.15 | 0.25 | 0 |
|  | locomotion pPC3 | 0.05 | 0.04 | 1.33 | 0.19 | 0 |
| Akaike weight: MC | ln(mass) | 0.2 | 0.15 | 1.36 | 0.18 | 0 |
|  | bill pPC1 | 0.12 | 0.15 | 0.81 | 0.42 | 0 |
|  | bill pPC2 | -0.02 | 0.14 | -0.12 | 0.9 | 0 |
|  | bill pPC3 | 0.2 | 0.12 | 1.64 | 0.11 | 0 |
|  | locomotion pPC1 | 0.16 | 0.15 | 1.05 | 0.3 | 0 |
|  | locomotion pPC2 | 0.05 | 0.13 | 0.35 | 0.73 | 0 |
|  | **locomotion pPC3** | **0.41** | **0.13** | **3.29** | **0.002** | **0** |
| relative support for a | ln(mass) | 0.08 | 0.14 | 0.53 | 0.6 | 0 |
| model with competition | bill pPC1 | 0.17 | 0.13 | 1.3 | 0.2 | 0 |
|  | bill pPC2 | 0 | 0.15 | 0 | 1 | 0 |
|  | bill pPC3 | 0.09 | 0.13 | 0.7 | 0.48 | 0 |
|  | locomotion pPC1 | 0.09 | 0.13 | 0.7 | 0.49 | 0 |
|  | locomotion pPC2 | -0.05 | 0.15 | -0.31 | 0.76 | 0 |
|  | locomotion pPC3 | 0.24 | 0.14 | 1.74 | 0.09 | 0 |

**S5 Table.** Intercept-only PGLS models fit to indices of support for two regimes models for each trait (for cases where N $\geq$ 50; *n* = 59). The index of relative support for any two-regime model was calculated using max(two regime Akaike weight)/(max(two regime Akaike weight)+max(single regime Akaike weight)); other, model specific indices were calculated using max(two regime Akaike weight for specified model)/ (max(two regime Akaike weight for specified model) + max(single regime Akaike weight for specified model)). For each model, this index was transformed by subtracting 0.5 such that negative estimates indicate support for a single-regime model and positive values equal support for a two-regime model. Values indicated in bold are those that are significant after controlling for multiple testing (α = 0.05/7). For all significant cases, the single-regime version of the model was supported over the two-regime version. λ indicates the maximum likelihood estimate of the phylogenetic signal.

| **response variable** | **trait** | **estimate** | **s.e.** | ***t*-value** | ***p*-value** | ***λ*** |
| --- | --- | --- | --- | --- | --- | --- |
| relative support for | ln(mass) | 0.04 | 0.04 | 0.89 | 0.38 | 0.11 |
| any two-regime model | bill pPC1 | 0.03 | 0.03 | 1.10 | 0.28 | 0 |
|  | bill pPC2 | -0.01 | 0.02 | -0.47 | 0.64 | 0 |
|  | bill pPC3 | 0.00 | 0.02 | 0.04 | 0.97 | 0 |
|  | locomotion pPC1 | 0.01 | 0.02 | 0.40 | 0.69 | 0 |
|  | locomotion pPC2 | 0.07 | 0.03 | 2.53 | 0.01 | 0 |
|  | locomotion pPC3 | 0.04 | 0.03 | 1.34 | 0.18 | 0 |
| relative support for | ln(mass) | 0.00 | 0.02 | 0.17 | 0.86 | 0 |
| two-regime BM model | bill pPC1 | 0.00 | 0.03 | 0.18 | 0.86 | 0 |
|  | bill pPC2 | -0.02 | 0.02 | -0.75 | 0.46 | 0 |
|  | **bill pPC3** | **-0.06** | **0.02** | **-3.84** | **0.0003** | **0** |
|  | locomotion pPC1 | -0.02 | 0.02 | -0.89 | 0.38 | 0 |
|  | locomotion pPC2 | -0.02 | 0.02 | -1.01 | 0.31 | 0 |
|  | locomotion pPC3 | 0.01 | 0.02 | 0.39 | 0.70 | 0 |
| relative support for | **ln(mass)** | **-0.10** | **0.02** | **-5.82** | **< 0.0001** | **0** |
| two-regime OU model | bill pPC1 | -0.06 | 0.02 | -2.75 | 0.01 | 0 |
|  | **bill pPC2** | **-0.09** | **0.02** | **-4.81** | **< 0.0001** | **0** |
|  | **bill pPC3** | **-0.07** | **0.02** | **-3.66** | **0.0006** | **0** |
|  | **locomotion pPC1** | **-0.08** | **0.02** | **-3.95** | **0.0002** | **0** |
|  | locomotion pPC2 | -0.02 | 0.03 | -0.83 | 0.41 | 0 |
|  | locomotion pPC3 | 0.06 | 0.03 | 2.1 | 0.04 | 0 |
| relative support for | ln(mass) | 0.04 | 0.06 | 0.61 | 0.54 | 0.38 |
| two-regime EB model | bill pPC1 | 0.02 | 0.03 | 0.77 | 0.44 | 0 |
|  | bill pPC2 | 0.03 | 0.03 | 0.95 | 0.34 | 0 |
|  | bill pPC3 | -0.02 | 0.02 | -0.65 | 0.52 | 0 |
|  | locomotion pPC1 | 0.00 | 0.02 | 0.00 | 1.00 | 0 |
|  | locomotion pPC2 | 0.03 | 0.03 | 1.17 | 0.25 | 0 |
|  | locomotion pPC3 | 0.01 | 0.03 | 0.23 | 0.82 | 0 |
| relative support for | ln(mass) | -0.04 | 0.08 | -0.43 | 0.67 | 0.72 |
| two-regime DD_exp_ model | bill pPC1 | 0.00 | 0.02 | 0.19 | 0.85 | 0 |
|  | bill pPC2 | 0.01 | 0.06 | 0.23 | 0.82 | 0.42 |
|  | bill pPC3 | -0.03 | 0.02 | -1.26 | 0.21 | 0 |
|  | locomotion pPC1 | -0.01 | 0.03 | -0.29 | 0.78 | 0 |
|  | locomotion pPC2 | 0.03 | 0.03 | 1.24 | 0.22 | 0 |
|  | locomotion pPC3 | -0.03 | 0.02 | -1.04 | 0.30 | 0 |
| relative support for | ln(mass) | -0.08 | 0.03 | -2.73 | 0.01 | 0 |
| two-regime DD_lin_ model | bill pPC1 | -0.01 | 0.04 | -0.25 | 0.80 | 0 |
|  | bill pPC2 | 0.09 | 0.11 | 0.76 | 0.45 | 0.55 |
|  | bill pPC3 | -0.06 | 0.04 | -1.54 | 0.13 | 0 |
|  | locomotion pPC1 | -0.03 | 0.03 | -1.01 | 0.32 | 0 |
|  | locomotion pPC2 | -0.09 | 0.03 | -2.68 | 0.01 | 0 |
|  | **locomotion pPC3** | **-0.12** | **0.04** | **-3.48** | **0.001** | **0** |
| relative support for | ln(mass) | -0.08 | 0.07 | -1.20 | 0.24 | 0.65 |
| two-regime MC model | bill pPC1 | -0.08 | 0.07 | -1.28 | 0.21 | 0.52 |
|  | **bill pPC2** | **-0.16** | **0.02** | **-7.67** | **< 0.0001** | **0** |
|  | **bill pPC3** | **-0.19** | **0.02** | **-10.98** | **< 0.0001** | **0** |
|  | locomotion pPC1 | -0.12 | 0.05 | -2.30 | 0.03 | 0.41 |
|  | **locomotion pPC2** | **-0.16** | **0.02** | **-7.96** | **< 0.0001** | **0** |
|  | **locomotion pPC3** | **-0.18** | **0.02** | **-8.02** | **< 0.0001** | **0** |

**S6 Table.** PGLS models comparing the observed latitudinal distribution (measured as the proportion of lineages with individuals that breed in tropical regions) of clade-by-trait level fits with the mean maximum likelihood estimates (across fits conducted on a bank of stochastic maps of ancestral biogeography) of the strength of species interactions in single-regime models incorporating competition. All comparisons were conducted on clades with $\geq$ 50 species (*n* = 66). Note: one outlier was removed from the exponential diversity dependence analysis of bill pPC2. Values indicated in bold are those that are significant after controlling for multiple testing (α = 0.05/7). λ indicates the maximum likelihood estimate of the phylogenetic signal.

| **model (parameter)** | **trait** | **estimate** | **std. error** | **t-value** | **p-value** | **λ** |
| --- | --- | --- | --- | --- | --- | --- |
| DDexp (slope) | ln(mass) | -0.01 | 0.05 | -0.28 | 0.78 | 1 |
|  | bill pPC1 | 0 | 0.03 | 0.04 | 0.97 | 0.42 |
|  | bill pPC2 | -1.67 | 1.19 | -1.41 | 0.16 | 0.35 |
|  | bill pPC3 | -0.87 | 1.27 | -0.68 | 0.5 | 0 |
|  | locomotion pPC1 | -0.18 | 0.5 | -0.36 | 0.72 | 1 |
|  | locomotion pPC2 | -1.97 | 1.16 | -1.7 | 0.09 | 0 |
|  | locomotion pPC3 | 0.38 | 0.26 | 1.48 | 0.14 | 0 |
| DDlin (slope) | ln(mass) | -9.92E-05 | 9.64E-05 | -1.03 | 0.31 | 0.56 |
|  | bill pPC1 | -5.18E-05 | 5.58E-05 | -0.93 | 0.36 | 0.53 |
|  | bill pPC2 | -3.51E-06 | 1.11E-05 | -0.32 | 0.75 | 0 |
|  | bill pPC3 | -4.26E-07 | 4.07E-06 | -0.1 | 0.92 | 0 |
|  | locomotion pPC1 | -2.82E-05 | 3.77E-05 | -0.75 | 0.46 | 0 |
|  | locomotion pPC2 | 8.97E-06 | 9.19E-06 | 0.98 | 0.33 | 0.47 |
|  | locomotion pPC3 | 2.86E-06 | 3.38E-06 | 0.85 | 0.4 | 0 |
| MC (S) | ln(mass) | -0.06 | 0.03 | -2.15 | 0.04 | 0 |
|  | bill pPC1 | -0.06 | 0.04 | -1.52 | 0.13 | 0 |
|  | bill pPC2 | -0.18 | 0.1 | -1.9 | 0.06 | 0.58 |
|  | bill pPC3 | -0.12 | 0.06 | -2.01 | 0.05 | 0 |
|  | locomotion pPC1 | -0.06 | 0.03 | -1.87 | 0.07 | 0 |
|  | locomotion pPC2 | 0.01 | 0.04 | 0.26 | 0.8 | 0 |
|  | **locomotion pPC3** | **-0.1** | **0.02** | **-4.1** | **0.0001** | **0** |

**S7 Table.** Intercept-only PGLS models linear regressions fit to tropical/temperate comparisons of maximum likelihood parameter estimates of the strength of species interactions in two-regime models (for cases where N$\geq$50) (*n* = 59) for each trait. For each evolutionary model (a: MC, b: DD_exp_, c: DD_lin_), the mean (across fits conducted on a bank of stochastic maps of ancestral biogeography and stochastic maps of breeding range) of the log-transformed ratio of the absolute value of parameter estimates for tropical taxa to that of temperate taxa (ln(|par_tropical|/|par_temperate|)) was the response variable in the intercept-only PGLS model. Negative estimates, therefore, indicate that the impact of competition is estimated to be higher in temperate regions, whereas positive estimates indicate that competition is higher in the tropics. Values indicated in bold are those that are significant after controlling for multiple testing (α = 0.05/7). λ indicates the maximum likelihood estimate of the phylogenetic signal.

| **response variable** | **model term** | **estimate** | **s.e.** | ***t*-value** | ***p*-value** | **λ** |
| --- | --- | --- | --- | --- | --- | --- |
| MC | ln(mass) | -1.68 | 0.82 | -2.04 | 0.05 | 0 |
| (ln(\|S_tropical_\|/ \|S_temperate_\|)) | bill pPC1 | -0.93 | 0.93 | -0.99 | 0.32 | 0 |
|  | **bill pPC2** | **-2.40** | **0.80** | **-3.00** | **0.004** | **0** |
|  | bill pPC3 | -0.88 | 0.77 | -1.14 | 0.26 | 0 |
|  | locomotion pPC1 | -0.71 | 0.97 | -0.73 | 0.47 | 0 |
|  | locomotion pPC2 | 1.98 | 0.96 | 2.07 | 0.04 | 0 |
|  | locomotion pPC3 | -1.90 | 0.90 | -2.11 | 0.04 | 0 |
| DD_exp_ | ln(mass) | -0.08 | 0.24 | -0.34 | 0.73 | 0 |
| (ln(\|r_tropical_\|/ \|r_temperate_\|)) | bill pPC1 | -0.10 | 0.27 | -0.35 | 0.72 | 0 |
|  | bill pPC2 | 0.11 | 0.30 | 0.36 | 0.72 | 0 |
|  | bill pPC3 | -0.48 | 0.31 | -1.53 | 0.13 | 0 |
|  | locomotion pPC1 | -0.45 | 0.27 | -1.70 | 0.09 | 0 |
|  | locomotion pPC2 | -0.40 | 0.22 | -1.79 | 0.08 | 0 |
|  | locomotion pPC3 | 0.09 | 0.26 | 0.33 | 0.74 | 0 |
| DD_lin_ | ln(mass) | -0.11 | 0.43 | -0.26 | 0.80 | 0 |
| (ln(\|b_tropical_\|/ \|b_temperate_\|)) | bill pPC1 | -0.51 | 0.87 | -0.59 | 0.56 | 0.19 |
|  | bill pPC2 | -1.11 | 0.75 | -1.49 | 0.14 | 0 |
|  | bill pPC3 | -0.77 | 0.80 | -0.97 | 0.34 | 0 |
|  | locomotion pPC1 | -0.28 | 0.50 | -0.55 | 0.58 | 0 |
|  | locomotion pPC2 | -0.83 | 0.68 | -1.21 | 0.23 | 0 |
|  | locomotion pPC3 | -1.56 | 0.77 | -2.03 | 0.05 | 0 |

**S8 Table.** Intercept-only PGLS models linear regressions fit to tropical/temperate comparisons of maximum likelihood parameter estimates of the strength of species interactions in two-regime models (for cases where N$\geq$100) (*n* = 34) for each trait. For each evolutionary model (a: MC, b: DD_exp_, c: DD_lin_), the mean (across fits conducted on a bank of stochastic maps of ancestral biogeography and stochastic maps of breeding range) of the log-transformed ratio of the absolute value of parameter estimates for tropical taxa to that of temperate taxa (ln(|par_tropical|/|par_temperate|)) was the response variable in the intercept-only PGLS model. Negative estimates, therefore, indicate that the impact of competition is estimated to be higher in temperate regions, whereas positive estimates indicate that competition is higher in the tropics. Values indicated in bold are those that are significant after controlling for multiple testing (α = 0.05/7). λ indicates the maximum likelihood estimate of the phylogenetic signal.

| **response variable** | **model term** | **estimate** | **s.e.** | ***t*-value** | ***p*-value** | **λ** |
| --- | --- | --- | --- | --- | --- | --- |
| MC | ln(mass) | -1.42 | 1.18 | -1.2 | 0.24 | 0 |
| (ln(\|S_tropical_\|/ \|S_temperate_\|)) | bill pPC1 | -0.52 | 1.25 | -0.42 | 0.68 | 0 |
|  | bill pPC2 | -1.89 | 0.91 | -2.08 | 0.05 | 0 |
|  | bill pPC3 | -1.07 | 1.08 | -0.99 | 0.33 | 0 |
|  | locomotion pPC1 | -0.67 | 1.33 | -0.5 | 0.62 | 0 |
|  | locomotion pPC2 | 2.94 | 1.3 | 2.27 | 0.03 | 0 |
|  | locomotion pPC3 | -1.01 | 1.33 | -0.76 | 0.45 | 0 |
| DD_exp_ | ln(mass) | -0.08 | 0.3 | -0.28 | 0.78 | 0 |
| (ln(\|r_tropical_\|/ \|r_temperate_\|)) | bill pPC1 | 0.23 | 0.34 | 0.66 | 0.51 | 0 |
|  | bill pPC2 | 0.22 | 0.36 | 0.6 | 0.55 | 0 |
|  | bill pPC3 | -0.4 | 0.35 | -1.13 | 0.27 | 0 |
|  | locomotion pPC1 | -0.22 | 0.33 | -0.68 | 0.5 | 0 |
|  | locomotion pPC2 | -0.46 | 0.26 | -1.78 | 0.08 | 0 |
|  | locomotion pPC3 | 0.21 | 0.35 | 0.6 | 0.55 | 0 |
| DD_lin_ | ln(mass) | -0.39 | 0.58 | -0.67 | 0.51 | 0 |
| (ln(\|b_tropical_\|/ \|b_temperate_\|)) | bill pPC1 | -0.05 | 0.56 | -0.08 | 0.93 | 0 |
|  | bill pPC2 | -1.12 | 0.96 | -1.16 | 0.25 | 0 |
|  | bill pPC3 | -0.85 | 1.1 | -0.77 | 0.44 | 0 |
|  | locomotion pPC1 | -0.11 | 0.66 | -0.17 | 0.87 | 0 |
|  | locomotion pPC2 | -0.81 | 0.91 | -0.89 | 0.38 | 0 |
|  | locomotion pPC3 | -1.47 | 1.06 | -1.39 | 0.17 | 0 |

**S9 Table.** Zero-intercept mixed-effect linear model with a random effect for clade identity fit to the proportion of lineages pairs in each clade that are sympatric in each continent.

| **response variable** | **model term** | **estimate** | **s.e.** | ***t*-value** |
| --- | --- | --- | --- | --- |
| Proportion of species pairs sympatric | W Palearctic | 0.74 | 0.04 | 19.82 |
|  | E Palearctic | 0.54 | 0.03 | 16.43 |
|  | W Nearctic | 0.70 | 0.04 | 19.74 |
|  | E Nearctic | 0.66 | 0.04 | 18.59 |
|  | Madagascar | 0.68 | 0.04 | 16.65 |
|  | Africa | 0.36 | 0.03 | 11.44 |
|  | South America | 0.34 | 0.03 | 10.66 |
|  | Central America | 0.41 | 0.03 | 13.2 |
|  | India | 0.63 | 0.03 | 19.59 |
|  | SE Asia | 0.40 | 0.03 | 12.89 |
|  | Oceania | 0.37 | 0.04 | 10.45 |

**S10 Table.** Intercept-only mixed-effect linear model with a random effect for clade identity fit to the proportion of lineages pairs that are sympatric in each clade.

| **response variable** | **model term** | **estimate** | **s.e.** | ***t*-value** |
| --- | --- | --- | --- | --- |
| Proportion of species pairs sympatric | intercept | 0.50 | 0.02 | 26.32 |

**S11 Table.** Linear model fit to the proportion of lineages pairs that are sympatric as a function of the absolute value of midpoint latitude for species pairs.

| **response variable** | **model term** | **estimate** | **s.e.** | ***z*-value** | ***p*-value** |
| --- | --- | --- | --- | --- | --- |
| Proportion of species pairs sympatric | intercept | -0.98 | 0.01 | -126.46 | < 0.001 |
|  | abs(midpoint latitude) | 0.02 | 0.0004 | 43.12 | < 0.001 |

**S12 Table.** Simulation parameters used in simulation study to explore the statistical power of two-regime MC models under realistic levels of sympatry. Values were chosen based on maximum likelihood estimates (MLEs) from single-regime MC models.

| **clade** | **simulation parameters** | | | |
| --- | --- | --- | --- | --- |
|  | *σ^2^* | *S*_tropical_ | *S*_temperate_ | S_tropical_:S_temperate_ |
| Cracidae.0  (MLE for single-regime MC:  *σ^2^* = 0.00464021,  *S* = -0.1269343) | 0.00464021 | -0.1269343 | -0.1269343 | 1:1 |
|  | 0.00464021 | -0.3808029 | -0.1269343 | 3:1 |
|  | 0.00464021 | -0.1904015 | -0.1269343 | 3:2 |
|  | 0.00464021 | -0.1269343 | -0.1904015 | 2:3 |
|  | 0.00464021 | -0.1269343 | -0.3808029 | 1:3 |
| Nectariniidae.0  (MLE for single-regime MC:  *σ^2^* = 0.00000676,  *S* = -0.1626235) | 0.00000676 | -0.1626235 | -0.1626235 | 1:1 |
|  | 0.00000676 | -0.4878705 | -0.1626235 | 3:1 |
|  | 0.00000676 | -0.2439352 | -0.1626235 | 3:2 |
|  | 0.00000676 | -0.1626235 | -0.2439352 | 2:3 |
|  | 0.00000676 | -0.1626235 | -0.4878705 | 1:3 |
| Picidae.1  (MLE for single-regime MC:  *σ^2^* = 0.00684987,  *S* = -0.0691007) | 0.00684987 | -0.0691007 | -0.0691007 | 1:1 |
|  | 0.00684987 | -0.207302 | -0.0691007 | 3:1 |
|  | 0.00684987 | -0.103651 | -0.0691007 | 3:2 |
|  | 0.00684987 | -0.0691007 | -0.103651 | 2:3 |
|  | 0.00684987 | -0.0691007 | -0.207302 | 1:3 |

**S13 Table.** PGLS analyses of maximum likelihood estimates of evolutionary rates in single-regime model fits (*n* = 135) as a function of the latitudinal distribution (measured as the proportion of lineages with individuals that breed in tropical regions). For diversity-dependent models, parameter estimates are the mean estimates across fits conducted on a bank of stochastic maps of ancestral biogeography. λ indicates the maximum likelihood estimate of the phylogenetic signal.

| **model (parameter)** | **trait** | **estimate** | **std. error** | ***t*-value** | ***p*-value** | **λ** |
| --- | --- | --- | --- | --- | --- | --- |
| BM (σ^2^) | ln(mass) | -0.0015 | 0.0024 | -0.64 | 0.52 | 0 |
|  | bill pPC1 | -0.0017 | 0.0014 | -1.18 | 0.24 | 0 |
|  | bill pPC2 | -0.00016 | 0.00028 | -0.56 | 0.58 | 0 |
|  | bill pPC3 | -8.58E-05 | 8.93E-05 | -0.96 | 0.34 | 0 |
|  | locomotion pPC1 | 0.0014 | 0.0013 | 1.07 | 0.29 | 0 |
|  | locomotion pPC2 | 0.00029 | 0.00035 | 0.81 | 0.42 | 0 |
|  | locomotion pPC3 | -5.71E-05 | 9.15E-05 | -0.62 | 0.53 | 0 |
| DDexp (maximum σ^2^ | ln(mass) | -0.0022 | 0.0025 | -0.86 | 0.39 | 0 |
| at the tips) | bill pPC1 | -0.0025 | 0.0015 | -1.71 | 0.09 | 0 |
|  | bill pPC2 | -0.00029 | 0.00026 | -1.11 | 0.27 | 0 |
|  | bill pPC3 | -0.00011 | 0.00013 | -0.84 | 0.4 | 0.86 |
|  | locomotion pPC1 | 0.0016 | 0.0018 | 0.92 | 0.36 | 0 |
|  | locomotion pPC2 | 0.00062 | 0.00052 | 1.19 | 0.24 | 0 |
|  | locomotion pPC3 | -0.00011 | 0.00012 | -0.94 | 0.35 | 0 |
| DDlin (maximum σ^2^ | ln(mass) | -0.0024 | 0.0025 | -0.98 | 0.33 | 0 |
| at the tips) | bill pPC1 | -0.0027 | 0.0015 | -1.79 | 0.08 | 0 |
|  | bill pPC2 | -0.00021 | 0.00027 | -0.78 | 0.44 | 0 |
|  | bill pPC3 | -0.00017 | 8.37E-05 | -1.97 | 0.05 | 0 |
|  | locomotion pPC1 | 0.00093 | 0.0016 | 0.6 | 0.55 | 0 |
|  | locomotion pPC2 | 0.00035 | 0.00046 | 0.76 | 0.45 | 0 |
|  | locomotion pPC3 | -0.00016 | 9.50E-05 | -1.71 | 0.09 | 0 |
| EB (σ^2^ at the tips) | ln(mass) | -0.0013 | 0.0021 | -0.63 | 0.53 | 0 |
|  | bill pPC1 | -0.00073 | 0.0010 | -0.72 | 0.47 | 0 |
|  | bill pPC2 | -0.00015 | 0.00019 | -0.82 | 0.42 | 0 |
|  | bill pPC3 | -5.20E-05 | 7.57E-05 | -0.69 | 0.49 | 0 |
|  | locomotion pPC1 | 0.0013 | 0.0012 | 1.11 | 0.27 | 0 |
|  | locomotion pPC2 | 0.00032 | 0.00035 | 0.92 | 0.36 | 0 |
|  | locomotion pPC3 | -5.11E-05 | 8.95E-05 | -0.57 | 0.57 | 0 |

**S14 Table.** Intercept-only PGLS models fit to the difference between tropical and temperate maximum likelihood parameter estimates of evolutionary rates in two-regime models, fit separately for each trait (*n* = 71 for ln.mass and *n* = 70 for other traits). For DD models, the rate parameter was calculated as the mean comparisons between parameter estimates across fits conducted on a bank of stochastic maps of ancestral biogeography and stochastic maps of breeding range. Note: one outlier was removed from the linear diversity dependence analysis of locomotion pPC2 as it was > 2 orders of magnitude larger than the next largest value. Values indicated in bold are those that are significant after controlling for multiple testing (α = 0.05/7). λ indicates the maximum likelihood estimate of the phylogenetic signal.

| **response variable** | **model term** | **estimate** | **s.e.** | ***t*-value** | ***p*-value** | **λ** |
| --- | --- | --- | --- | --- | --- | --- |
| BM (σ^2^_tropical_ - σ^2^_temperate_) | ln(mass) | -0.0037 | 0.0029 | -1.26 | 0.21 | 0.48 |
|  | bill pPC1 | 1.5E-05 | 0.00052 | 0.03 | 0.98 | 0 |
|  | bill pPC2 | -0.00023 | 8.0E-05 | -2.88 | 0.01 | 0 |
|  | bill pPC3 | 2.40E-05 | 4.0E-05 | 0.60 | 0.55 | 0 |
|  | locomotion pPC1 | 0.00017 | 0.00042 | 0.40 | 0.69 | 0 |
|  | locomotion pPC2 | 0.0001 | 0.00016 | 0.67 | 0.51 | 0 |
|  | **locomotion pPC3** | **-0.00014** | **4.30E-05** | **-3.29** | **0.002** | **0** |
| EB (σ^2^_tropical_ - σ^2^_temperate_) | ln(mass) | -0.0014 | 0.00086 | -1.66 | 0.10 | 0 |
|  | bill pPC1 | 0.00013 | 0.00048 | 0.27 | 0.79 | 0 |
|  | bill pPC2 | -0.00015 | 5.6E-05 | -2.64 | 0.01 | 0 |
|  | bill pPC3 | 0.00011 | 0.0001 | 1.09 | 0.28 | 0.48 |
|  | locomotion pPC1 | 0.00022 | 0.00035 | 0.64 | 0.53 | 0 |
|  | locomotion pPC2 | 0.00012 | 0.00014 | 0.84 | 0.40 | 0 |
|  | **locomotion pPC3** | **-0.00011** | **3.4E-05** | **-3.23** | **0.002** | **0** |
| DD_exp_ (σ^2^_tropical_ - σ^2^_temperate_) | ln(mass) | -0.0023 | 0.0014 | -1.59 | 0.12 | 0 |
|  | bill pPC1 | -0.002 | 0.0093 | -0.22 | 0.83 | 1 |
|  | **bill pPC2** | **-0.00032** | **7.5E-05** | **-4.22** | **< 0.0001** | **0** |
|  | bill pPC3 | 0.00023 | 0.00026 | 0.88 | 0.38 | 0.64 |
|  | locomotion pPC1 | -0.00075 | 0.0015 | -0.52 | 0.61 | 0 |
|  | locomotion pPC2 | -0.00015 | 0.001 | -0.14 | 0.89 | 0.5 |
|  | locomotion pPC3 | -0.00043 | 0.00031 | -1.42 | 0.16 | 0.5 |
| DD_lin_ (σ^2^_tropical_ - σ^2^_temperate_) | ln(mass) | -0.0053 | 0.003 | -1.80 | 0.08 | 0.29 |
|  | bill pPC1 | -0.0018 | 0.0017 | -1.06 | 0.30 | 0.26 |
|  | **bill pPC2** | **-0.00023** | **6.4E-05** | **-3.59** | **0.0006** | **0** |
|  | bill pPC3 | -6.8E-06 | 4.3E-05 | -0.16 | 0.88 | 0 |
|  | locomotion pPC1 | -0.00044 | 0.00052 | -0.85 | 0.40 | 0 |
|  | locomotion pPC2 | 7.8E-05 | 0.00024 | 0.33 | 0.74 | 0 |
|  | locomotion pPC3 | -0.00011 | 8.4E-05 | -1.34 | 0.19 | 0 |

**S15 Table.** PGLS models comparing the observed latitudinal distribution (measured as the proportion of lineages with individuals that breed in tropical regions) of clade-by-trait level fits (*n* = 135) with the log-transformed error (calculated as the sum of the maximum likelihood estimated error parameter and the clade-level mean squared standard error) in single-regime Brownian motion models. Values indicated in bold are those that are significant after controlling for multiple testing (α = 0.05/7). λ indicates the maximum likelihood estimate of the phylogenetic signal.

| **trait** | **intercept** | **estimate** | **std. error** | ***t*-value** | ***p*-value** | **λ** |
| --- | --- | --- | --- | --- | --- | --- |
| **ln(mass)** | **-3.92** | **-1.41** | **0.5** | **-2.83** | **0.0054** | **0** |
| bill pPC1 | -4.49 | -0.38 | 0.46 | -0.82 | 0.41 | 0.37 |
| bill pPC2 | -5.55 | -0.29 | 0.33 | -0.86 | 0.39 | 0.46 |
| bill pPC3 | -6.38 | 0.24 | 0.28 | 0.83 | 0.41 | 0.55 |
| locomotion pPC1 | -4.88 | -0.53 | 0.43 | -1.22 | 0.23 | 0 |
| locomotion pPC2 | -5.45 | -0.49 | 0.35 | -1.39 | 0.17 | 0.4 |
| locomotion pPC3 | -6.99 | 0.04 | 0.31 | 0.13 | 0.9 | 0 |

**S16 Table.** PGLS models comparing the observed latitudinal distribution (measured as the proportion of lineages with individuals that breed in tropical regions) of clade-by-trait level fits (*n* = 135) with the maximum likelihood parameter estimates of evolutionary rates in single-regime Brownian motion models that do not account for observational error. Values indicated in bold are those that are significant after controlling for multiple testing (α = 0.05/7). λ indicates the maximum likelihood estimate of the phylogenetic signal.

| **model (parameter)** | **trait** | **estimate** | **std. error** | ***t*-value** | ***p*-value** | **λ** |
| --- | --- | --- | --- | --- | --- | --- |
| BM (σ^2^) | **ln(mass)** | **-0.012** | **0.0035** | **-3.38** | **0.0009** | **0.95** |
|  | bill pPC1 | -0.0064 | 0.0032 | -1.99 | 0.05 | 0 |
|  | bill pPC2 | -0.00087 | 0.00047 | -1.86 | 0.06 | 0.47 |
|  | bill pPC3 | -0.00024 | 0.00028 | -0.88 | 0.38 | 0.73 |
|  | locomotion pPC1 | -0.00065 | 0.0024 | -0.27 | 0.79 | 0 |
|  | locomotion pPC2 | -0.00022 | 0.00061 | -0.36 | 0.72 | 0 |
|  | **locomotion pPC3** | **-0.00075** | **0.00021** | **-3.54** | **0.0006** | **1** |

**S17 Table.** Intercept only PGLS models fit to the mean difference (across stochastic maps of tropical and temperate living) in MLE estimates of tropical and temperate rates (from two-rate BM models that do not account for observational error) (*n* = 71 for log-transformed body mass, 70 for other traits). Values indicated in bold are those that are significant after controlling for multiple testing (α = 0.05/7). λ indicates the maximum likelihood estimate of the phylogenetic signal.

| **response variable** | **model term** | **estimate** | **s.e.** | ***t*-value** | ***p*-value** | **λ** |
| --- | --- | --- | --- | --- | --- | --- |
| BM (σ^2^_tropical_ - σ^2^_temperate_) | **ln(mass)** | **-0.0039** | **0.0012** | **-3.15** | **0.002** | **0** |
|  | bill pPC1 | -0.0026 | 0.0016 | -1.61 | 0.11 | 0 |
|  | bill pPC2 | -0.00067 | 0.00024 | -2.76 | 0.01 | 0 |
|  | bill pPC3 | 4.1E-05 | 9.1E-05 | 0.45 | 0.65 | 0 |
|  | locomotion pPC1 | -0.0016 | 0.0013 | -1.21 | 0.23 | 0 |
|  | locomotion pPC2 | -0.00062 | 0.00032 | -1.96 | 0.05 | 0 |
|  | **locomotion pPC3** | **-0.00027** | **7.9E-05** | **-3.38** | **0.001** | **0** |

**S18 Table.** The factors predicting which clades support models with competition, as revealed by Phylogenetic Generalised Linear Mixed Models (PGLMMs) fit to single-regime clade-by-trait fits (*n* = 924) with a categorical variable indicating (a) that the matching competition was the modal best fit model (i.e., the most common best fit model across fits conducted on a bank of stochastic maps of ancestral biogeography) (*n* = 166) or (b) that the exponential diversity dependent model was the model best fit model (*n* = 66). The influence of the phylogeny was estimated from the random effect component of the PGLMM—the phylogenetic intraclass correlation coefficient is analogous to the λ parameter (often referred to as ‘phylogenetic signal’) estimated from phylogenetic generalized least squares models [20]. To facilitate parameter exploration, we rescaled all predictor variables using *z*-transformations. We used an uninformative, inverse Wishart distribution as a prior for the random effects, a flat prior for the fixed effects, and fixed the residual variance at 1 [21]. To fit the models, we ran an MCMC chain for at least 5 x 10^5^ generations, recording model results every 100 generations and ignoring the first 5 x 10^3^ generations as burn-in. We fit each model four times and merged the four chains after verifying convergence both visually and using Gelman-Rubin diagnostics in the R-package coda [22,23]. Estimates and credibility intervals are therefore calculated from the pooled posterior distributions. The pMCMC (an MCMC derived *p*-value calculated as two times the proportion of estimates in either the positive or negative portion (whichever is smaller) of the posterior distribution) is presented from one chain.

| **response variable** | **model term** | **median estimate** | **(95% CI)** | ***p*MCMC** |
| --- | --- | --- | --- | --- |
| (a) MC best fit == 1 | (intercept) | -1.68 | (-3.38, 0.09) | 0.06 |
| *λ* = 0.51 | clade age | -0.02 | (-0.45, 0.40) | 0.94 |
| (95% CI = 0.32, 0.67) | **clade richness** | **0.80** | **(0.42 , 1.23)** | **< 0.001** |
|  | **max prop. species coexisting** | **0.76** | **(0.31 , 1.26)** | **0.002** |
|  | prop. species strongly territorial | 0.21 | (-0.21, 0.64) | 0.32 |
|  | **prop. species single-strata habitats** | **-0.55** | **(-1.10, 0.07)** | **0.02** |
|  | diet diversity | 0.06 | (-0.35, 0.46) | 0.77 |
| (b) DD_exp_ best fit == 1 | **(intercept)** | **-3.41** | **(-4.11, -2.70)** | **< 0.001** |
| *λ* = 0.06 | clade age | -0.09 | (-0.44, 0.24) | 0.61 |
| (95% CI = 0.0002, 0.34) | **clade richness** | **0.51** | **(0.20, 0.84)** | **0.002** |
|  | **max prop. species coexisting** | **-0.71** | **(-1.11, -0.32)** | **< 0.001** |
|  | prop. species strongly territorial | 0.15 | (-0.21, 0.49) | 0.42 |
|  | prop. species single-strata habitats | -0.37 | (-0.83, 0.03) | 0.07 |
|  | diet diversity | -0.08 | (-0.42, 0.24) | 0.60 |

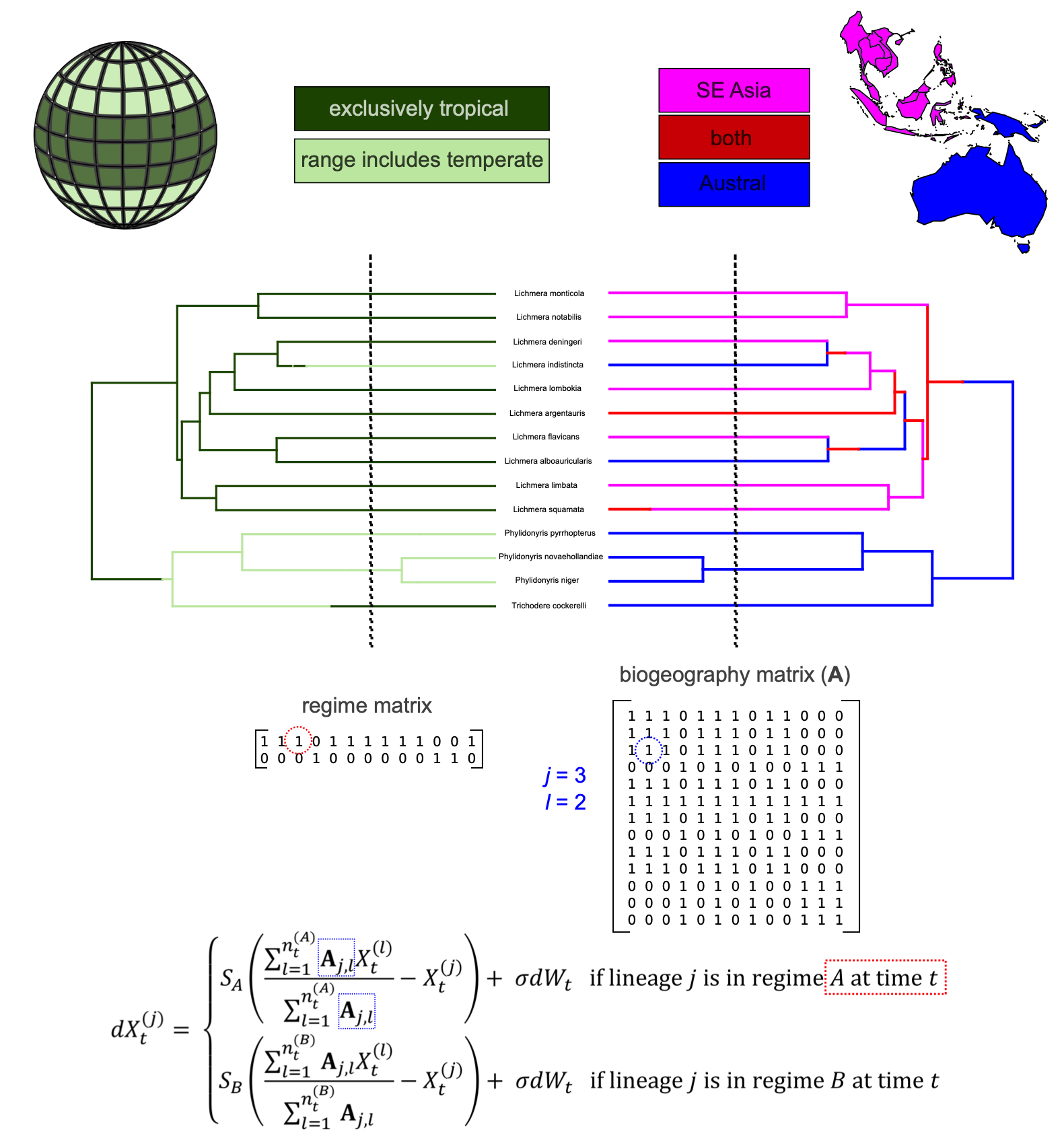

S1 Figure. Illustration of our model-fitting approach for clade-level model fits with different strengths of competition in tropical and temperate regions. We combine a matrix of the presence or absence of each lineage in tropical/temperate regions (‘regime matrix’) with a matrix of biogeography (denoted ‘**A’**) to identify the competitive regime of each lineage and the identity of other lineages with which the focal lineage is able to interact with. Blue and red colours in the lower panel denote correspondence between the formula and the biogeography matrix (**A**) and the regime matrix, respectively.

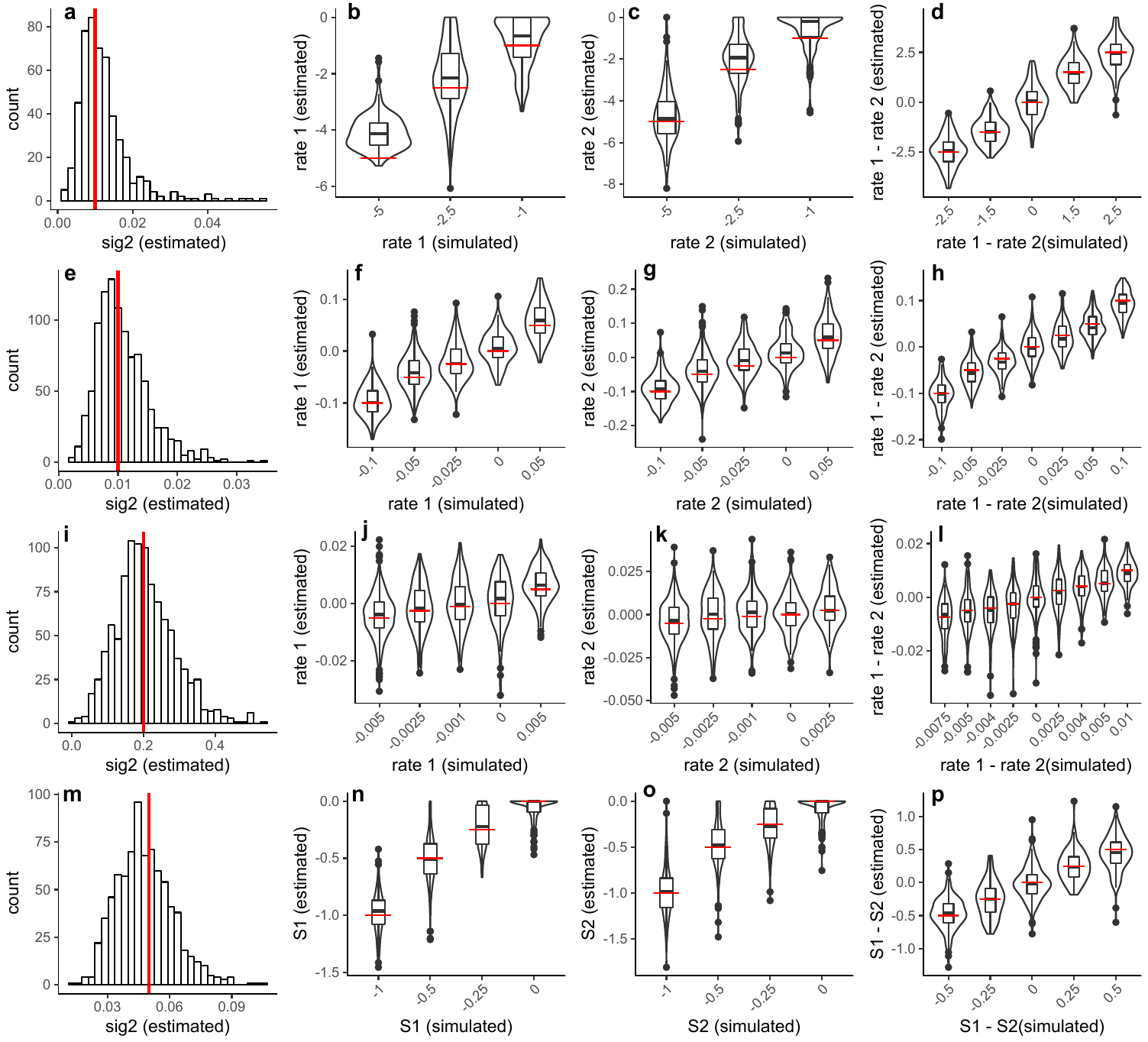

S2 Figure. Results of the simulation study demonstrate the maximum likelihood optimisation returns reliable parameter estimates in two-regime models. **a-d.** exponential time-dependent model **e-h.** exponential diversity-dependent model, **i-l.** linear diversity-dependent model, and **m-p.** matching competition model. In all plots, the red lines denote the parameters used to generate the simulated data.

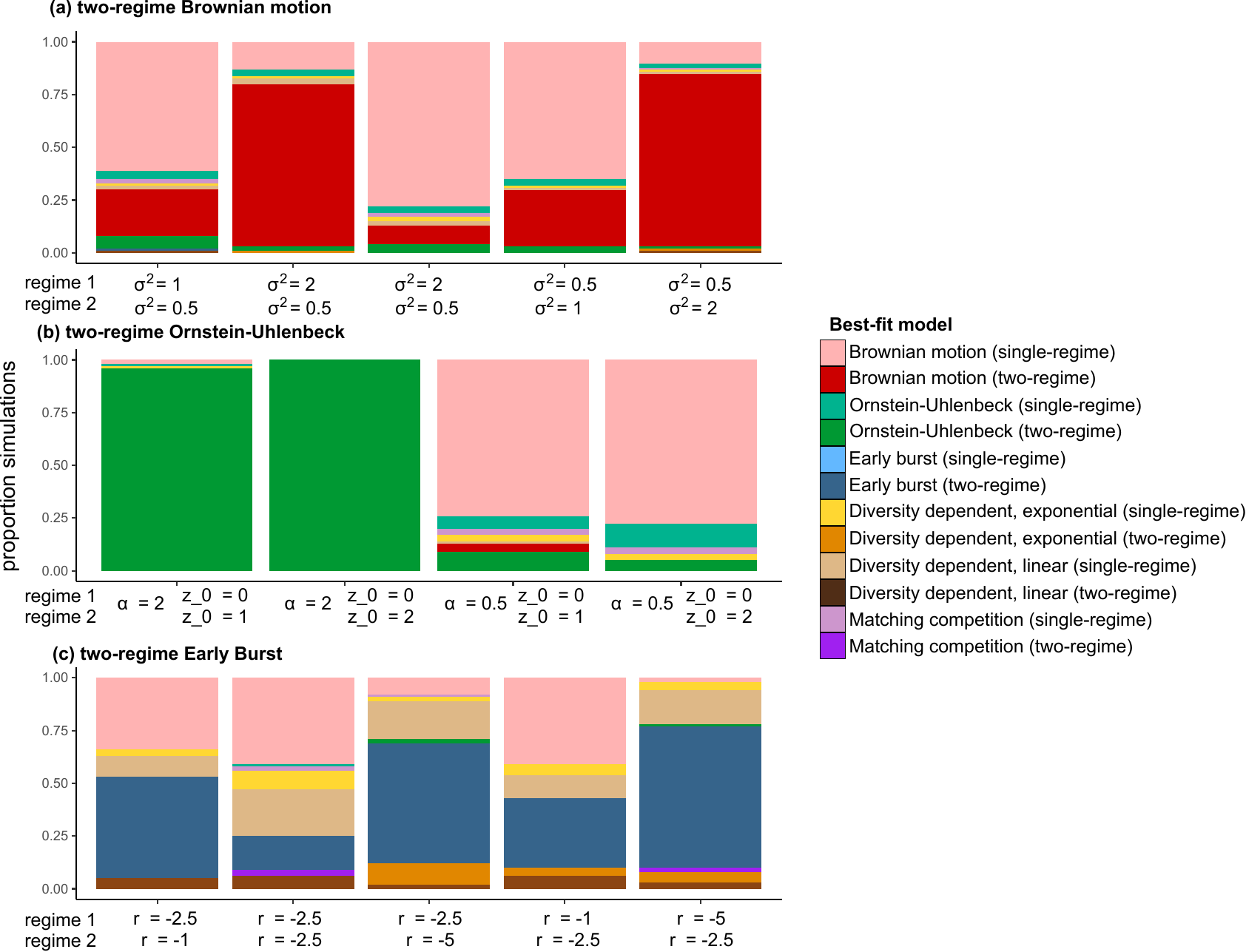

S3 Figure. Results of model selection depicting best fitting models for data simulated under (a) two-regime Brownian motion, (b) two-regime Ornstein-Uhlenbeck, and (c) two-regime Early Burst models across a range of parameter values.

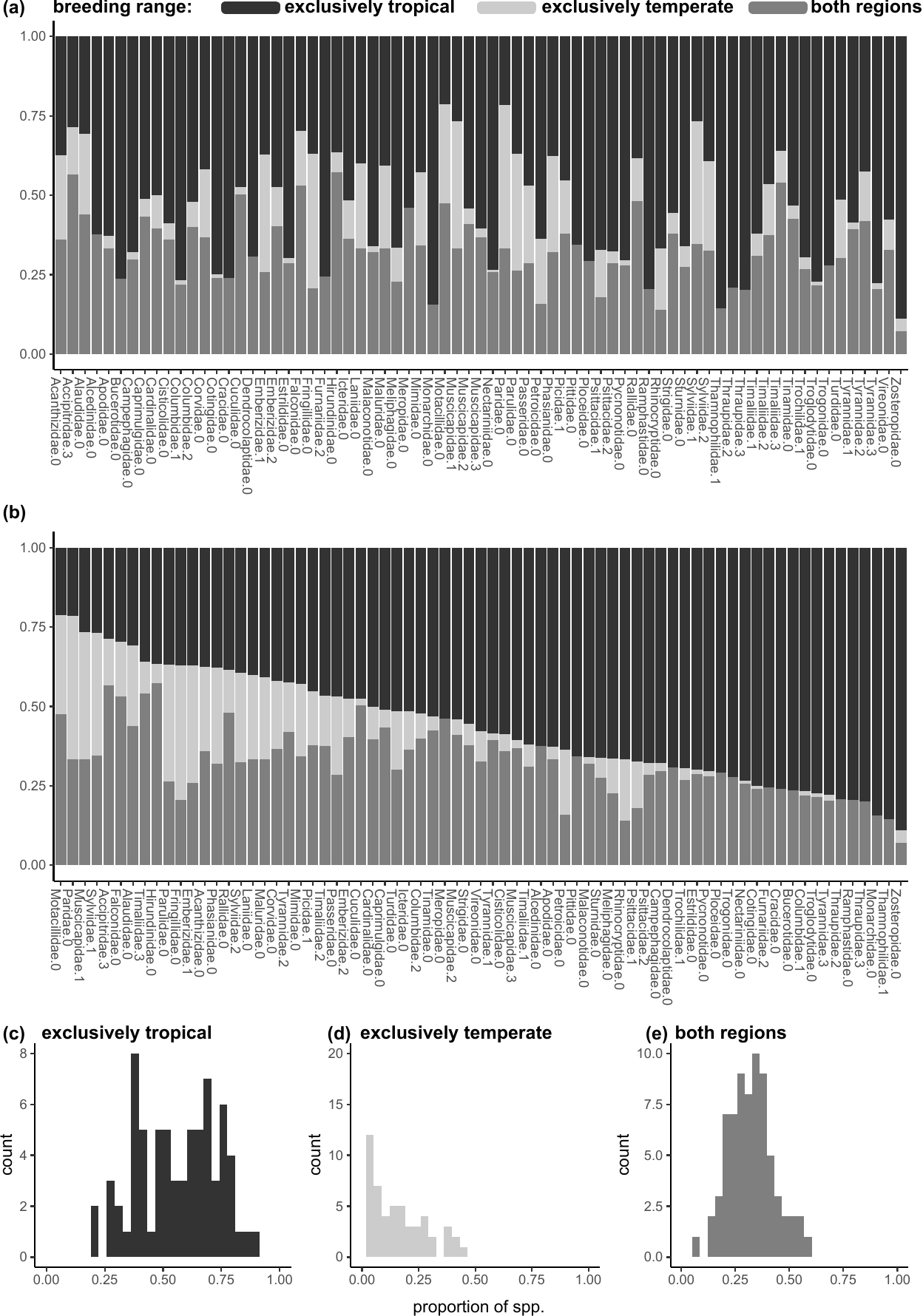

S4 Figure. Clade-level distributions of tropical, temperate, and widespread breeding (a) sorted by clade name, (b) sorted by proportion of exclusively tropical breeding species, and (c-d) presented as separate histograms. The number following the family name indicates the subclade within that family (see Methods).

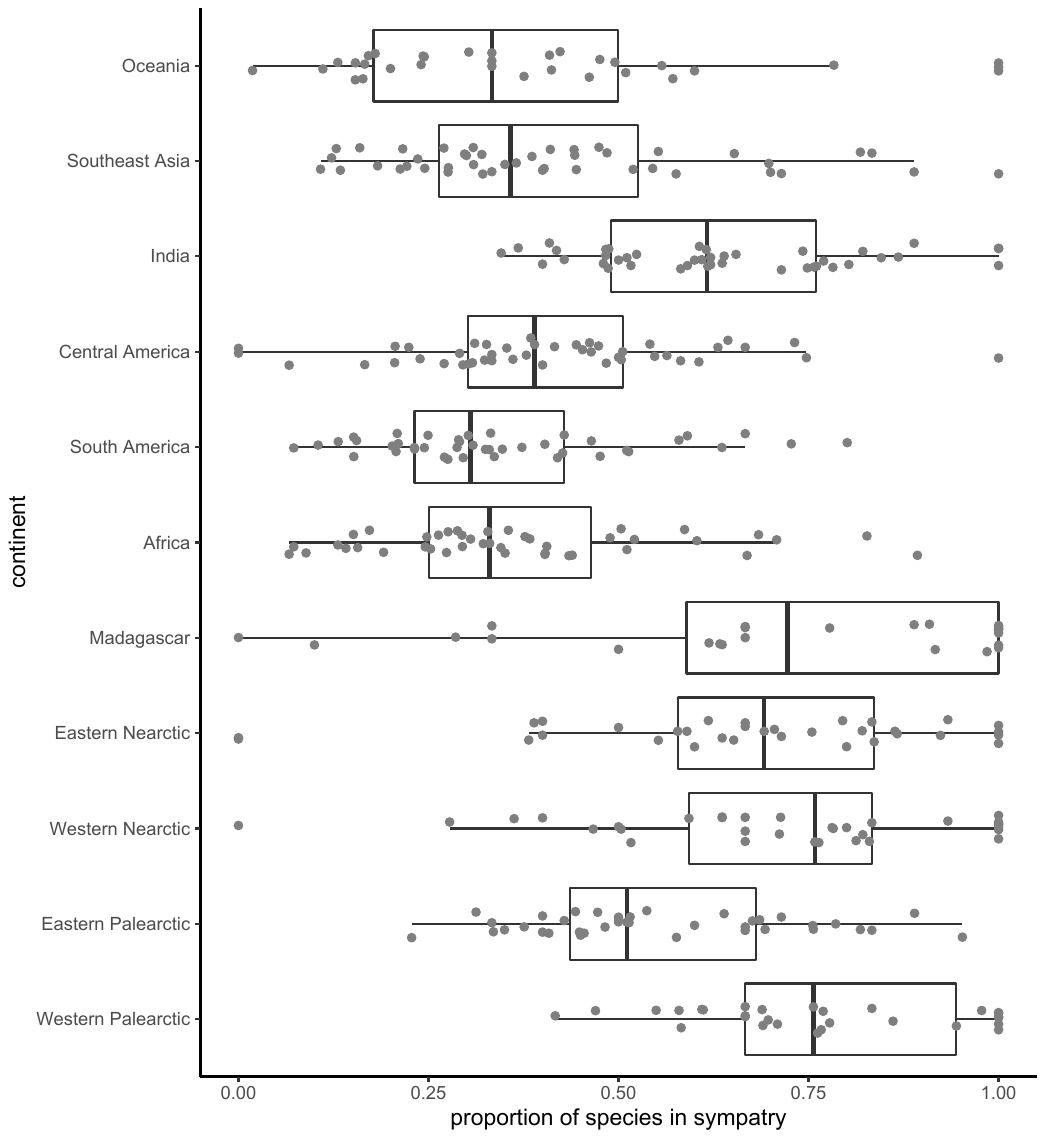

S5 Figure. Continental variation in the proportion of species that cooccur in sympatry (defined as 20% range overlap).

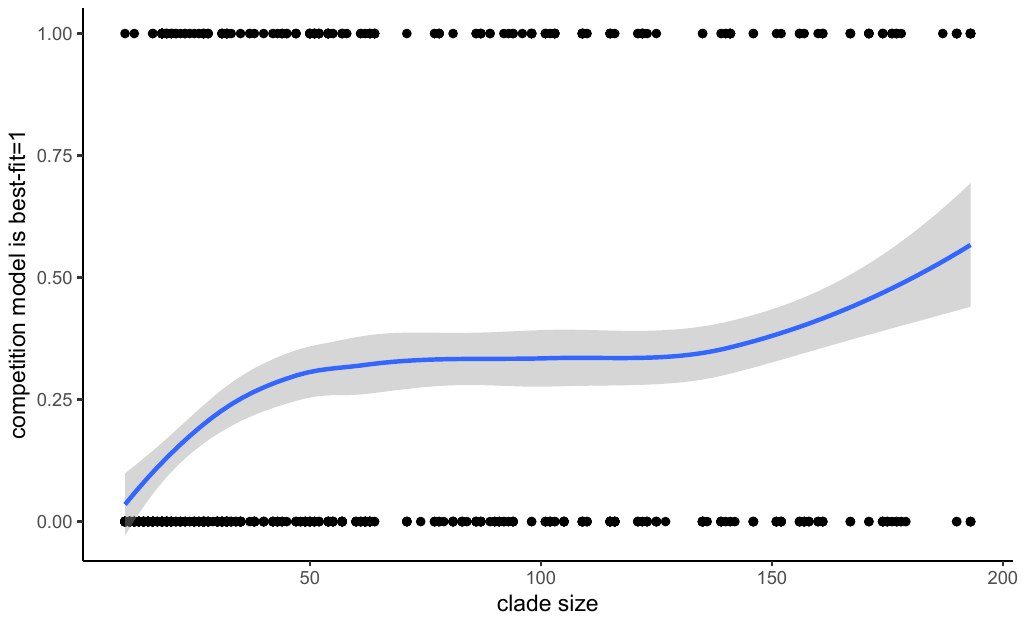

S6 Figure. Clade size impacts the probability that a model incorporating competition is the modal best-fit model (i.e., the most common best fit model across fits conducted on a bank of stochastic maps of ancestral biogeography and stochastic maps of breeding range).

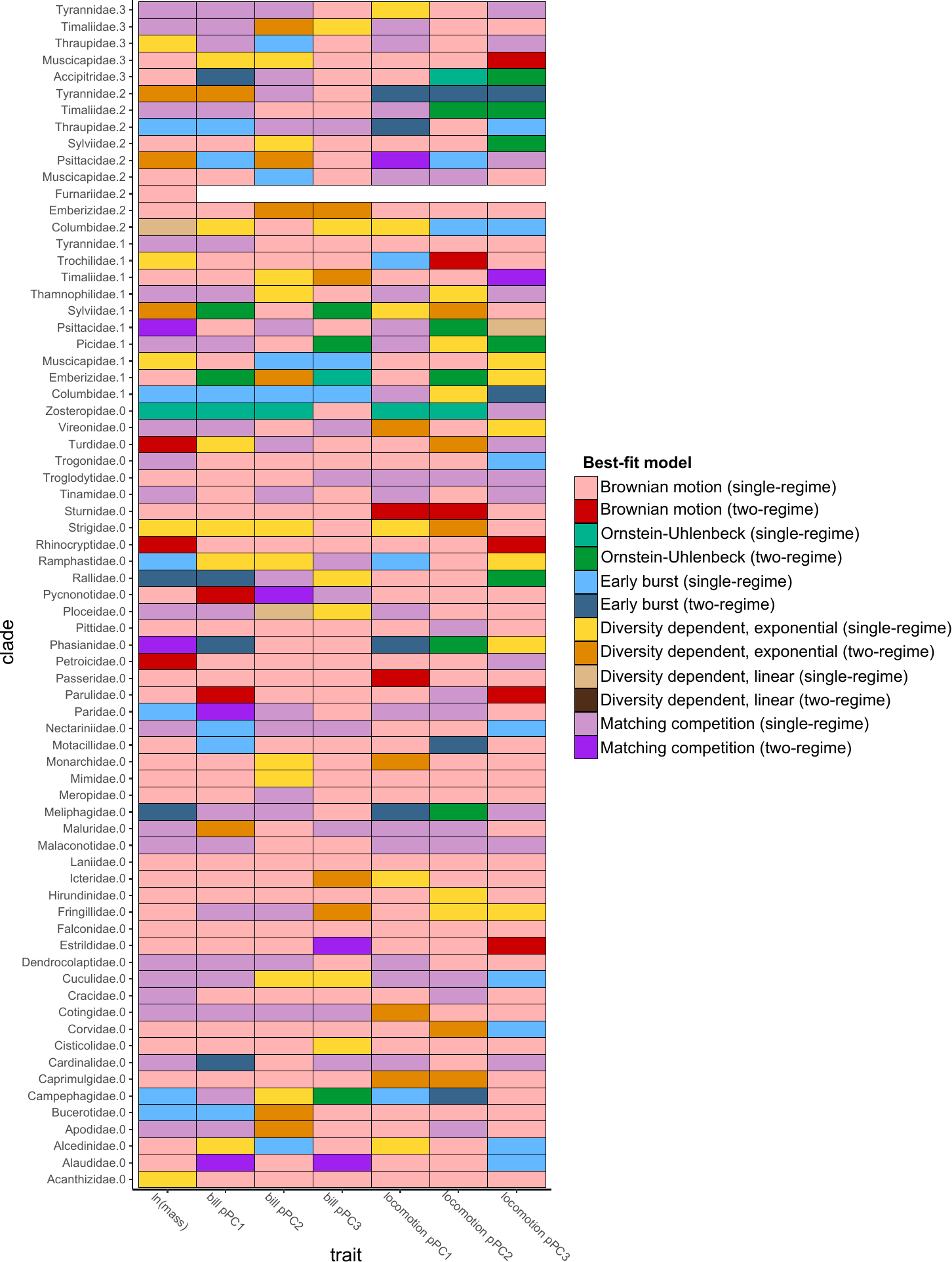
 S7 Figure. Best-fit models for each clade-by-trait combination shows that single-regime models generally outperform two-regime models, though some clades (e.g., Meliphagidae, Phasianidae) do tend to support models with latitude across several traits. Shown is the modal best-fit model (i.e., the most common best fit model across fits conducted on a bank of stochastic maps of ancestral biogeography across fits conducted on a bank of stochastic maps of ancestral biogeography and stochastic maps of breeding range). The number following the family name indicates the subclade within that family (see Methods).

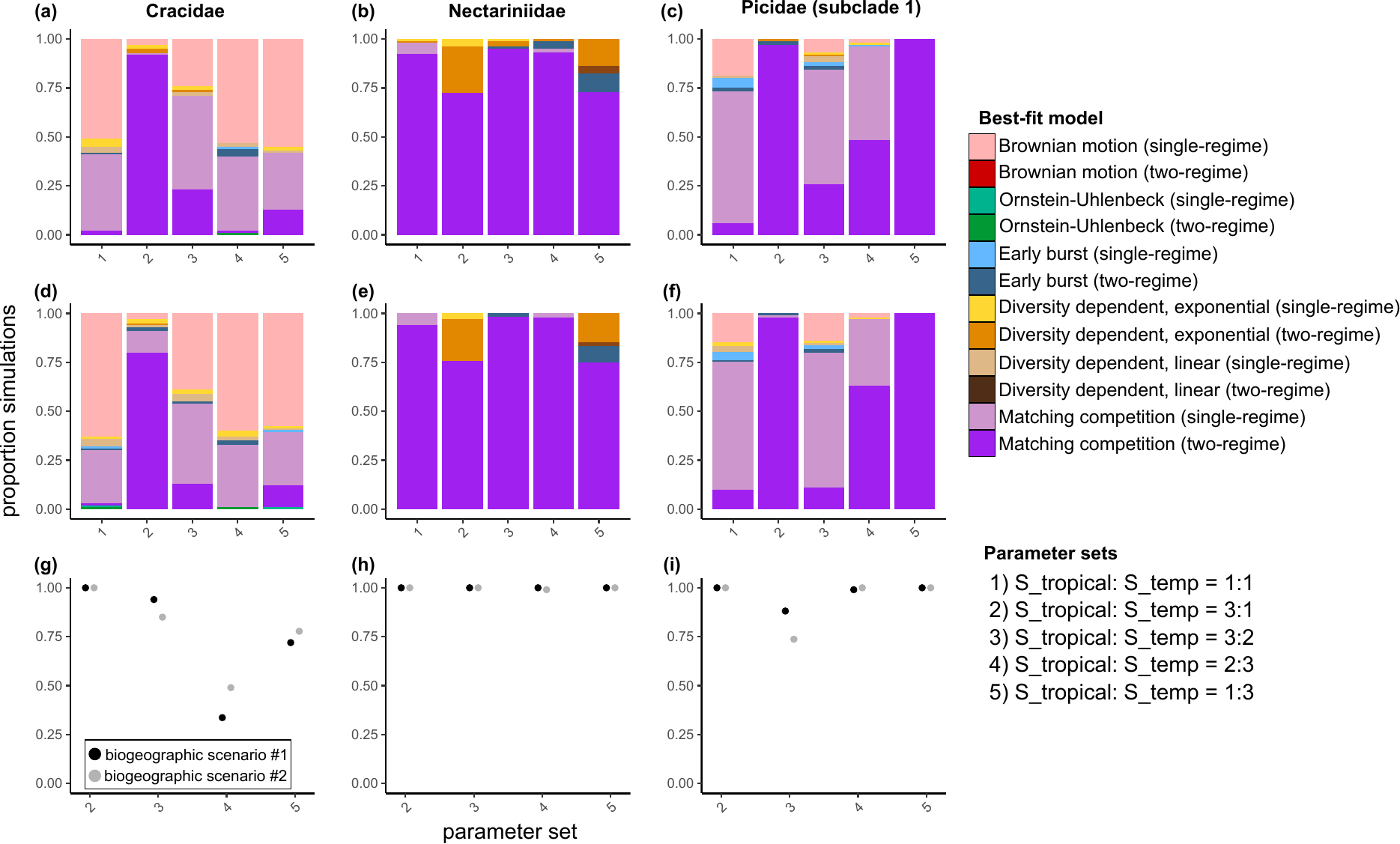

S8 Figure. Results from simulation analyses exploring the impact of assuming continental level sympatry for three clades. (a-c) Best-fit models for data generated under downsampled biogeographic scenario #1 (i.e., 50% of both tropical and temperate lineages set to allopatric at a continental scale). (d-f) Best-fit models for data generated under downsampled biogeographic scenario #2 (i.e., 50% of temperate lineages and 66.6% of tropical lineages set to allopatric at a continental scale). (g-i) The proportion of simulations for which maximum likelihood estimates of the ratio of competition from the two-regime MC model (i.e., ln(|S_tropical_|/|S_temperate_|) ) correctly identify the direction of the difference in the strength of competition.

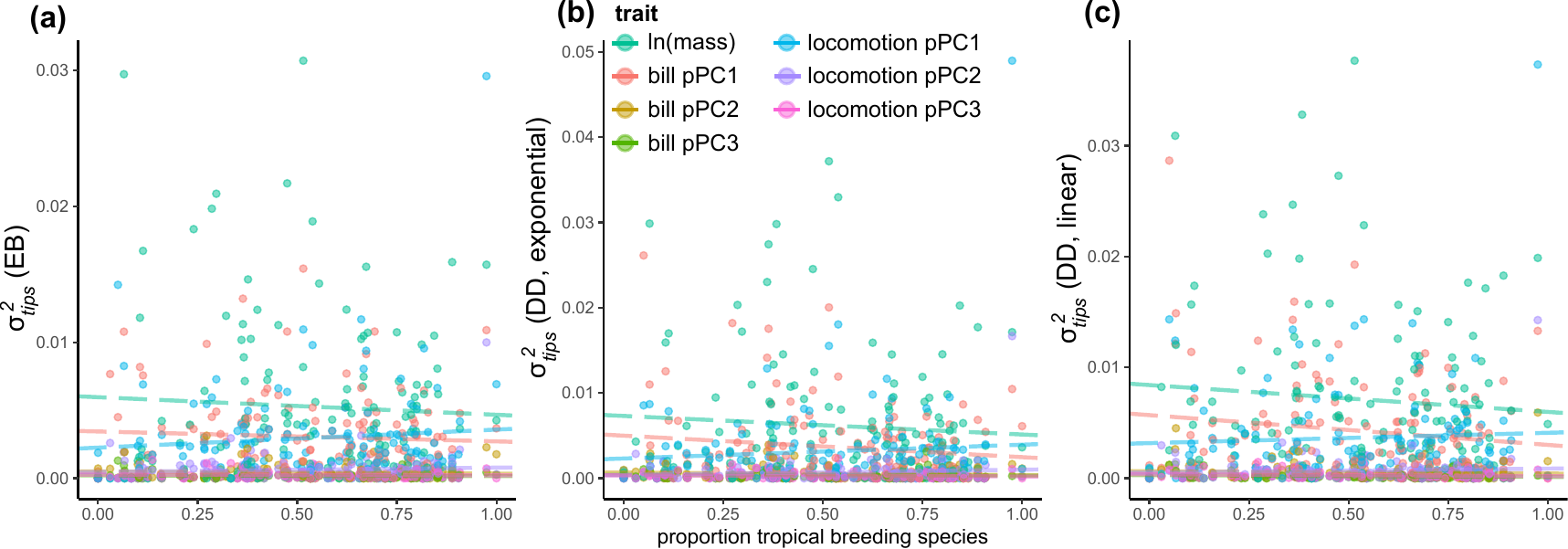

S9 Figure. Evolutionary rates in other single-regime models (a: EB, b: DDexp, c: DDlin) do not vary as a function of the proportion of lineages that breed in the tropics. For diversity-dependent models, parameter estimates are the mean estimates across fits conducted on a bank of stochastic maps of ancestral biogeography.

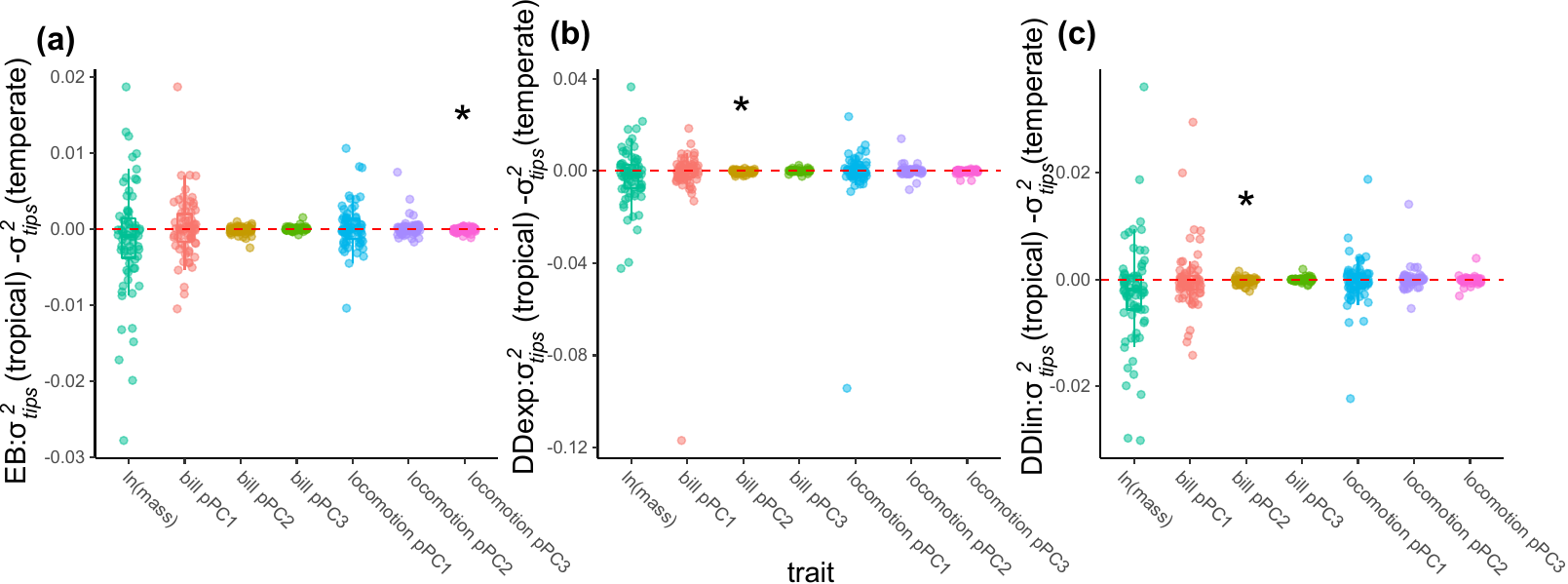

S10 Figure. Differences between rates estimated separately on tropical and temperate taxa in two-regime models (a: EB, b:DDexp, c: DDlin). Shown are the mean comparisons between parameter estimates across fits conducted on a bank of stochastic maps of ancestral biogeography and stochastic maps of breeding range (i.e., tropical or temperate). Asterisks indicate statistical significance.

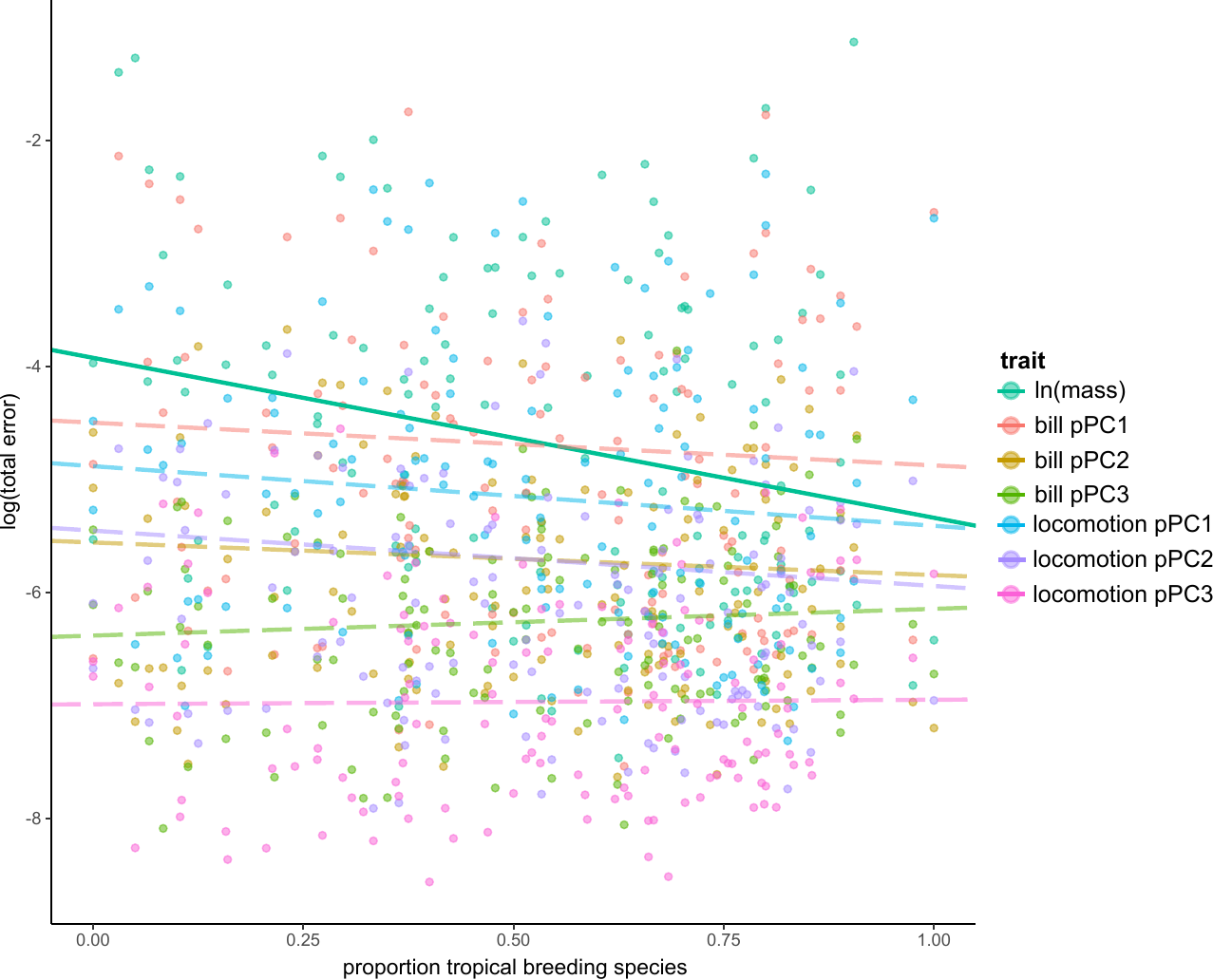

S11 Figure. The relationship between the total error (calculated as the log-transformed sum of the maximum likelihood estimated nuisance error parameter from single-regime Brownian motion models and the clade-level mean squared standard error) and the proportion of tropical breeding lineages in a clade is negative for body mass, but not for other traits. Solid lines represent statistically significant relationships (Table S9).

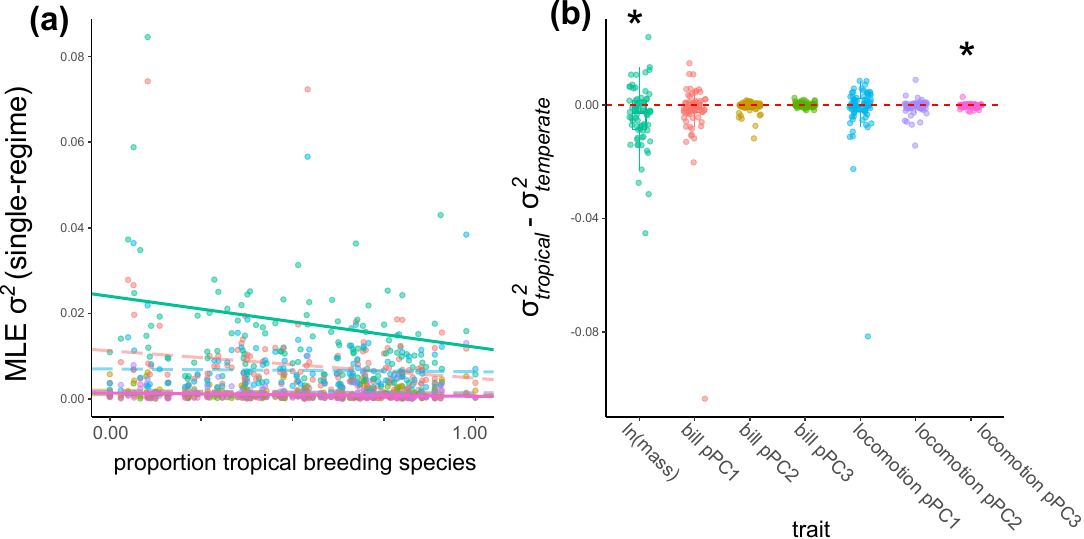

S12 Figure. Brownian motion models of trait evolution fit at a clade level when not accounting for observational error reveal a more pronounced relationship between rate and latitude for several traits **a.** There is a negative relationship between the proportion of taxa in a clade that breed in the tropics and the estimated rate of trait evolution from single-rate Brownian motion models for body mass and locomotion pPC3, but not other traits. Colour of points indicate trait (as in panel b). **b.** Differences between rates estimated separately on tropical and temperate taxa in two-rate Brownian motion models are biased toward faster rates in temperate regions for body mass and locomotion pPC3, but not other traits. Shown are the mean comparisons between parameter estimates across fits conducted on a bank of stochastic maps of ancestral biogeography and stochastic maps of breeding range (i.e., tropical or temperate).

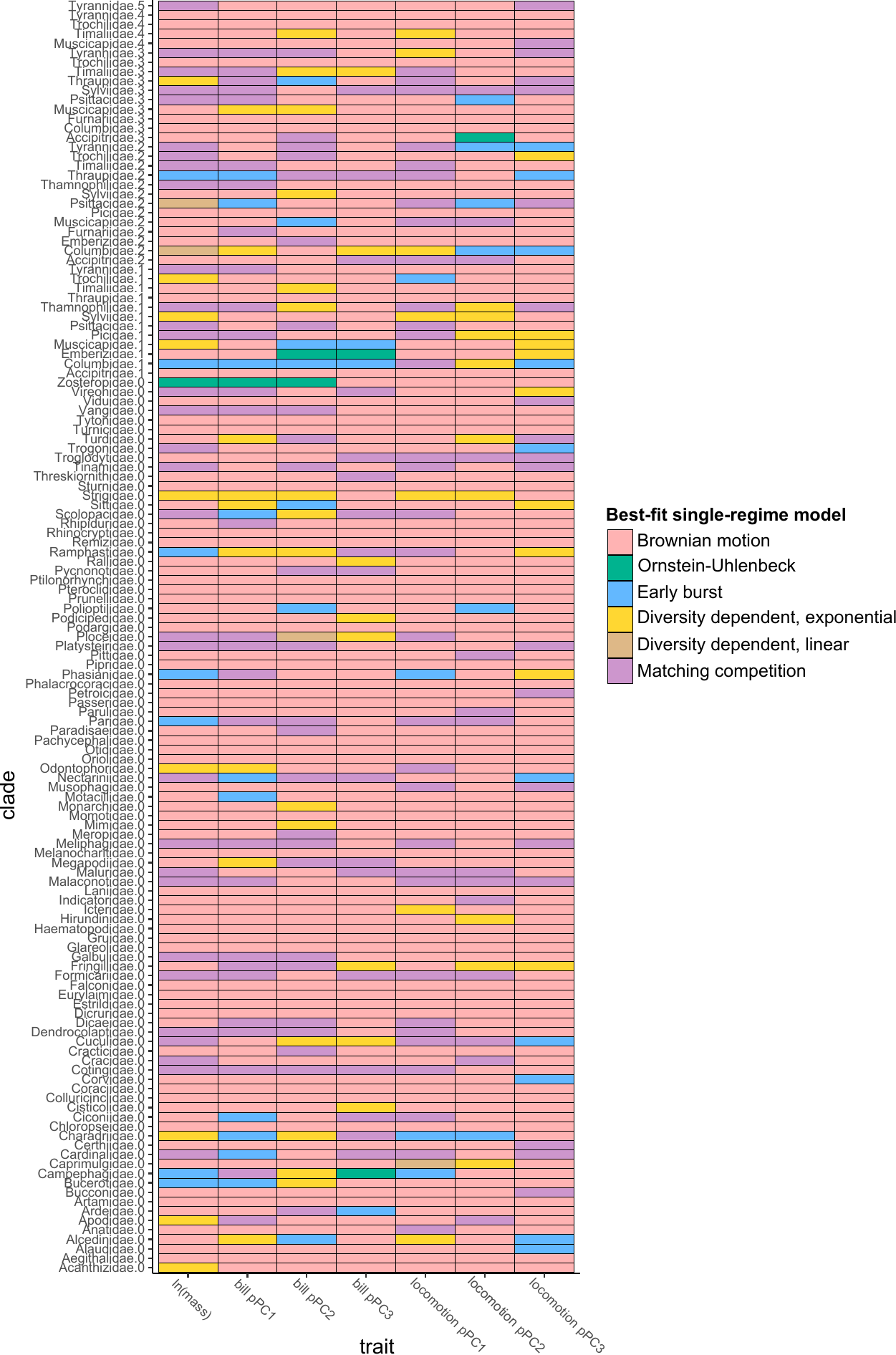

S13 Figure. Best-fit ‘single-regime’ models for each clade-by-trait combination show that, while Brownian motion is most often the best model, several clades show evidence of matching competition (e.g., Cotingidae, Formicariidae, Malaconotidae, and Paridae) or diversity dependence (e.g., Strigidae, Fringillidae, Columbidae subclade 2) acting on several traits. Shown is the modal best-fit model across fits conducted on a bank of stochastic maps of ancestral biogeography. The number following the family name indicates the subclade within that family (see Methods).

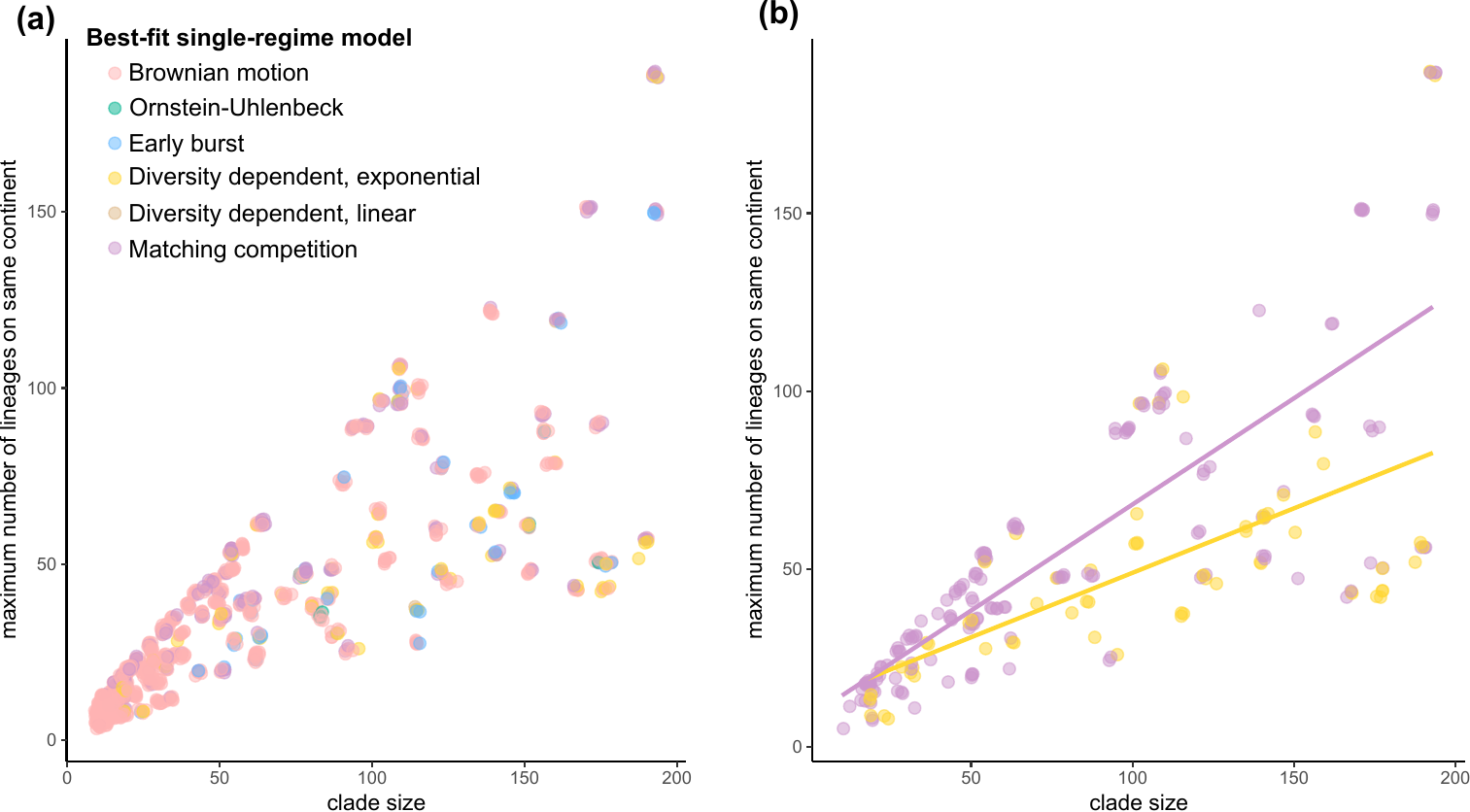

S14 Figure. Best-fit single-regime models (modal best fit across fits conducted on a bank of stochastic maps of ancestral biogeography), plotted as a function of total clade size and the number of species in each clade that occur on the same continent. A) All models, B) Matching competition and exponential diversity-dependent models. Each point represents a clade-by-trait combination (i.e., each clade contributes a point for each of seven traits). In both panels, points are jittered slightly to aid visualization.
